# Clustering Deviation Index (CDI): A robust and accurate unsupervised measure for evaluating scRNA-seq data clustering

**DOI:** 10.1101/2022.01.03.474840

**Authors:** Jiyuan Fang, Cliburn Chan, Kouros Owzar, Liuyang Wang, Diyuan Qin, Qi-Jing Li, Jichun Xie

## Abstract

Single-cell RNA-sequencing (scRNA-seq) technology allows us to explore cellular heterogeneity in the transcriptome. Because most scRNA-seq data analyses begin with cell clustering, its accuracy considerably impacts the validity of downstream analyses. Although many clustering methods have been developed, few tools are available to evaluate the clustering “goodness-of-fit” to the scRNA-seq data. In this paper, we propose a new Clustering Deviation Index (CDI) that measures the deviation of any clustering label set from the observed single-cell data. We conduct *in silico* and experimental scRNA-seq studies to show that CDI can select the optimal clustering label set. Particularly, CDI also informs the optimal tuning parameters for any given clustering method and the correct number of cluster components.

## 1 Introduction

Single-cell RNA-sequencing (scRNA-seq) quantifies the transcriptome of individual cells, allowing us to explore the biological heterogeneity among cells (Shapiro et al. 2013). Thus, scRNA-seq analysis usually begins with cell type clustering. Over the past five years, many methods have been developed or re-purposed for scRNA-seq clusterings, such as K-means, hierarchical clustering, RaceID (Grün et al. 2015), CIDR (Lin et al. 2017), SIMLR (Wang et al. 2017), SCANPY (Louvain algorithm) (Wolf et al. 2018), and Seurat (Louvain algorithm) (Stuart et al. 2019).

The outputs of these clustering methods are cell label sets that assign each cell to a cluster. Different methods usually yield different label sets. Even if we use a given clustering method, we still obtain different label sets by setting different tuning parameters. These different label sets lead to the challenge of choosing an “optimal” label set. To address the challenge, one approach is to apply a consensus method on these label sets to derive an ensemble label set. However, the ensemble label set is not guaranteed to reflect the underlying cell type structure better than any input label set. Furthermore, different consensus methods (Yang et al. 2019; Kiselev et al. 2017) often generate different ensemble label sets; thus, the challenge of choosing the optimal label set remains. Therefore, we need a reasonable evaluating index to score the “goodness-of-fit” or the deviation of each label set to the data. The evaluating index will help us select the optimal label set among the candidates.

In general, the evaluating indices can be divided into two categories. The first category consists of unsupervised indices. Calculating unsupervised indices does not depend on the knowledge of the actual label set; instad, the unsupervised indices usually use geometric or statistical properties to evaluate the quality of a label set. For example, the Calinski-Harabaz index (Caliński and Harabasz 1974) selects the label set with the minimum ratio between the within-cluster and between-cluster variances; similarly, the Silhouette coefficient (Rousseeuw 1987) and the Davies-Bouldin index (Davies and Bouldin 1979) selects the label set with the minimum ratio between the within-cluster and between-cluster distances. However, in the context of scRNA-seq clustering, we found that these methods often selected different label sets. None of their selected label sets matches the benchmark label set (Fig. 5).

The second category consists of supervised indices, whose accuracy is determined by the benchmark label set. Examples of supervised indices include the Adjusted Rand Index (ARI) (Hubert and Arabie 1985), the Normalized Mutual Information (Vinh et al. 2010), Fowlkes-Mallows scores (Fowlkes and Mallows 1983), and the weighted Rand Index and Mutual Information (Z Wu and H Wu 2020). One of the most commonly used supervised indices is ARI, a corrected-for-chance version of the Rand index that measures the agreement between two label sets: if they are similar, ARI is close to 1; otherwise close to 0. Because calculating supervised indices relies on the benchmark label set, the supervised indices are more accurate than unsupervised indices when the benchmark label set is accurate or close to the truth. Unfortunately, the benchmark label set is usually unavailable or challenging to generate. Moreover, even if it is available, it could be biased or incorrect because of outdated domain knowledge, leading to poor performances of supervised indices.

In this study, we developed a new unsupervised index, Clustering Deviation Index (CDI), to quantify the deviation of single cell data from the the data distribution based on the given label set. CDI is an unsupervised evaluation index whose calculation does not rely on the actual unobserved label set. However, its performance on scRNA-seq is consistent with ARI (Fig. 5). We applied CDI to multiple experimental scRNA-seq data sets and demonstrated that it successfully selected biologically meaningful clustering labels in each case. Because CDI is unsupervised, it is much more broadly applicable than supervised indices.

## 2 Results

### 2.1 Characterizing UMI count distributions of monoclonal cells

We built our model on a rigorous characterization of the statistical properties of the scRNA-seq data, CT26.WT, from a pure monoclonal population. We generated CT26.WT for a murine colon carcinoma cell line derived through monoclonal expansion to eliminate cell type heterogeneity and avoid experimental biases (Section Discussion). Then, we extracted the unique molecular identifier (UMI) count of each cell. UMI is barcoded for each transcript before amplification in many scRNA-seq protocols, leading to more accurate quantification of the transcript count (Klein et al. 2015; Zheng et al. 2017). Based on CT26.WT, we evaluated the “goodness-of-fit” of the following four families of gene-specific UMI count distributions to the actual gene-specific UMI count distributions. All four families consist of negative binomial (NB) distributions or zero-inflated NB distributions. The difference among these families lies in their dispersion and zero-inflation parameter modeling, and their mean parameter modeling is similar (Supplemental Note 1).

- *Gene-common NB*: negative binomial (NB) distributions with gene-common dispersion parameters;
- *Gene-common ZINB*: zero-inflated negative binomial (ZINB) distributions with gene-common dispersion parameters;
- *Gene-specific NB*: NB distributions with gene-specific dispersion parameters;
- *Gene-specific ZINB*: ZINB distributions with gene-specific dispersion parameters.

We used the Pearson’s chi-squared test (Chernoff and Lehmann 1954) to evaluate the “goodness-of-fit” of the distributions in the four distribution families to the actual gene-specific UMI count distributions in CT26.WT (Supplemental Fig. S2). The test rejected 34.3% poorly fitted genes for the gene-common NB family and 34.5% for the gene-common ZINB family. The rejection rates are high, indicating that these distribution families do not fit the actual UMI count distributions. In contrast, when applied to the gene-specific NB and ZINB families, the “goodness-of-fit” tests only rejected 9.7% and 6.1% poorly fitted genes, respectively. The rejection rates are not far from the preset type I error rate 5%, suggesting an overall good fit of these models.

The test results suggest that the well-fitted distribution family should include the gene-specific dispersion parameters, but including the zero-inflation parameters might not be necessary. To demonstrate the latter point, we split CT26.WT into two datasets - a half for training and the other half for testing. We fitted the unknown parameters in these four distribution families in the training dataset, estimated the zero UMI count proportions in the test dataset, and then compared the estimated and the observed zero UMI count proportions. The results (Fig. 1A) show that the gene-common NB and ZINB families underestimated the zero UMI count proportions in CT26.WT. In contrast, the gene-specific NB and ZINB families yielded reasonable and similar estimates of the zero UMI count proportions. Thus, adding additional ZINB parameters does not further improve the fitting. Hence, for the remainder of this study, we used the gene-specific-dispersion NB model without zero inflation.

**Figure 1.**
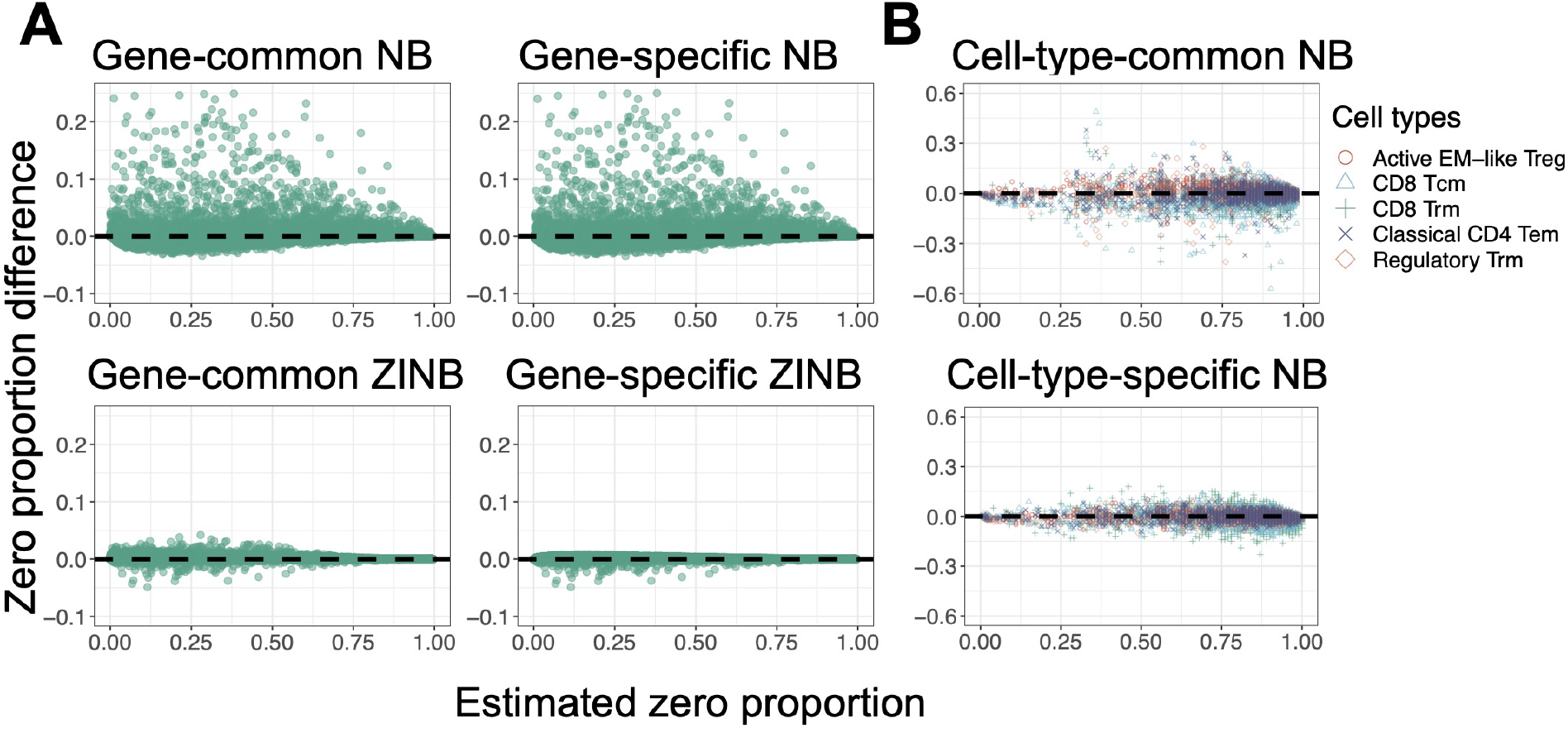
Zero UMI count proportion fitting under six models in monoclonal and polyclonal datasets. In both (*A*) and (*B*), cells in each type were randomly divided into two datasets: half for training and half for testing. The x-axis is the estimated zero UMI count probabilities based on the training dataset, and the y-axis is the difference between observed proportions in the test dataset and the estimated zero probabilities based on the training dataset. (*A*) The monoclonal CT26.WT dataset (*n* = 9, 621 cells). Each point represents a gene. (*B*) The polyclonal T-CELL dataset (n = 2, 989 cells). Each point represents a gene in a specific cell type, while the point colors and shapes represent five benchmark cell types (Christian et al. 2021). The CD8 Tcm cell type in this study is defined as *IL17RA^+^CD28^+^*.

### 2.2 Characterizing UMI count distributions of polyclonal cells

Polyclonal cell populations consist of cells from multiple cell types. To fit their UMI count distributions, we consider the following two NB distribution families. The difference between the two families lies in whether the mean and dispersion parameters are the same across cell types (Supplemental Note 1).

- *Cell-type-common NB*: NB distributions with the cell-type-common but gene-specific mean and dispersion parameters;
- *Cell-type-specific NB*: NB distributions with the cell-type-specific and gene-specific mean and dispersion parameters.

We used the cell-type-specific “goodness-of-fit” tests (Section 4.2) to check whether these two families fit the observed UMI counts well in the T-CELL dataset (Table 1). The tests rejected 34.27% of the genes for the cell-type-common NB family and 2.27% for the cell-type-specific NB family. Thus, adding cell-type-specific parameters substantially improved the fitting to the UMI count distributions.

**Table 1.** Dataset summary. More details on the experimental and simulated datasets are provided in Sections 4.5 and 4.6.

|  | Dataset | # cells | # genes | # cell types (subtypes) | Protocols | Reference |
| --- | --- | --- | --- | --- | --- | --- |
| Real datasets | CT26.WT | 9,621 | 11,710 | 1(-) | 10X v3 | - |
|  | T-CELL | 2,989 | 7,893 | 5(-) | 10X | Christian et al. 2021 |
|  | CORTEX | 7,390 | 12,887 | 8(33) | inDrop | Hrvatin et al. 2018 |
|  | RETINA | 26,830 | 13,118 | 6(18) | Drop-seq | Shekhar et al. 2016 |
| Simulated datasets | SD1 | 4,000 | 10,000 | 10(-) | - | - |
|  | SD2 | 4,200 | 10,000 | 4(-) | - | - |
|  | SD3 | 2,800 | 10,000 | 2(4) | - | - |
|  | SD4 | 3,000 | 4,887 | 5(-) | - | - |

The cell-type-specific NB family consists of more free-changing parameters, so we need to show that the improved fit does not stem from overfitting. Therefore, we did a 50:50 train/test split on the T-CELL dataset. We used the training dataset to fit the gene-specific distributions in each family, estimated the zero UMI count proportions in the test dataset, and then compared the estimated and the observed zero UMI count proportions in the test dataset. In the test dataset, we found that the cell-type-specific NB family still provides much better estimates to the zero UMI count proportions (Fig. 1B). We performed similar analyses on the CORTEX and RETINA datasets and observed similar results (Supplemental Fig. S5–S8).

### 2.3 CDI Overview

We developed CDI as an unsupervised index to evaluate the fitting of the observed UMI counts to the UMI count distribution based on the candidate label set. It calculates the negative penalized maximum log-likelihood of the selected feature genes based on the candidate label set. We have shown that the raw UMI counts follow gene-specific and cell-type-specific NB distributions given the actual cell-type labels. CDI is low if the candidate label set and the actual label set are similar; otherwise, CDI is high. The CDI calculation involves the following two steps (Fig. 2).

**Figure 2.**
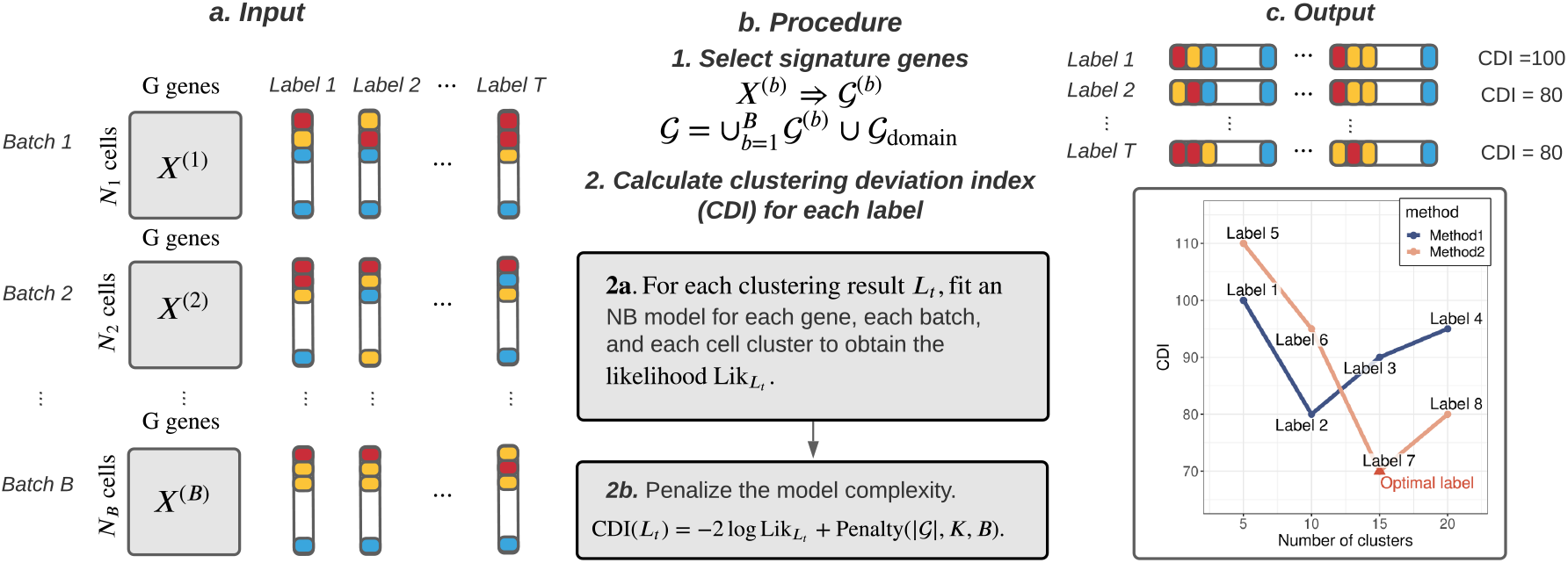
CDI calculation illustration. CDI inputs are *T* candidate label sets and the raw UMI count matrices from *B* batches. CDI contains two steps. CDI first selects the feature genes for each batch and takes their union. Other known feature genes can also be manually included. Then, CDI is calculated for all candidate label sets. CDI outputs are *T* CDI values corresponding to *T* candidate label sets. The label set with the lowest CDI is optimal.

1. **Feature gene selection.** Feature genes are those differentially expressed across cell types. Therefore, many scRNA-seq clustering methods rely on feature genes to cluster cells: selecting feature genes could substantially reduce data dimensions and possibly boost the signal in clustering. We also selected feature genes before calculating CDIs because of similar reasons. Many existing feature gene selection methods are available (Brennecke et al. 2013; Townes et al. 2019; Stuart et al. 2019). Here, we derive a new approach using a working dispersion score (WDS). WDS estimates the working dispersion for each gene. For single-batch datasets, we select genes with the largest average sample dispersion estimates as the feature genes. For multi-batch datasets, we rank genes in each batch by their average sample dispersion, combine rankings across batches by taking the minimum, and select top-ranked genes as the feature genes. Compared with other feature selection methods, our approach is derived from the parametric NB mixture model, capturing the fold change of mean parameters across cell types (Section S2.2). This property allows WDS to improve the performance of CDI, as described in Section 2.4.1, and Section 4.3.
2. **Optimal clustering label set selection.** If the candidate label set is close to the actual cell label set, the UMI count of each feature gene follows a gene-specific and cell-type-specific NB distribution. We calculated CDI as the sum of the negative penalized maximum log-likelihood for all the feature genes. The penalties are based on either the Akaike Information Criterion (AIC) or the Bayesian Information Criterion (BIC) (Akaike 1974; Schwarz et al. 1978; Konishi and Kitagawa 2008) (see details in Section 4.4). The choice of AIC or BIC depends on users’ scientific goals. Because BIC puts a higher penalty on model complexity, CDI with BIC (CDI-BIC) favors label sets with fewer clusters. Thus, we recommend using CDI-BIC to select the optimal label set on main cell types. Conversely, we recommend using CDI with AIC (CDI-AIC) to select the optimal subtype label set to depict the heterogeneity with a higher resolution.

### 2.4 Performance Evaluation

We evaluated the performance of WDS and CDI on four simulated datasets (SD1–SD4) and three experimental datasets (T-CELL, CORTEX, and RETINA). All datasets have the benchmark label sets. For the simulated datasets, the benchmark label sets are the actual cell label sets. For the experimental scRNA-seq datasets, the benchmark label sets were obtained by the multi-step process, including fluorescence-activated cell sorting (FACS), known feature gene checking, cell screening, and clustering; thus, these benchmark label sets may not be accurate but reflect our best knowledge of the cell types (see Section 4.6 for details).

#### 2.4.1 Performance of WDS in selecting feature genes for CDI

We compared WDS against another feature selection method, VST, the default for Seurat V3 (Stuart et al. 2019) and V4 (Hao et al. 2021). First, we selected the top 500 feature genes using WDS and VST, respectively. Second, we normalized the UMI counts of the selected feature genes by the log(max(count, 0.1)) transformation. Third, we calculated the top 50 principal components (PCs) of the normalized UMI counts. Finally, we plotted the two-dimensional uniform manifold approximation and projection (UMAP) (Becht et al. 2018) based on the top 50 PCs (Fig. 3).

**Figure 3.**
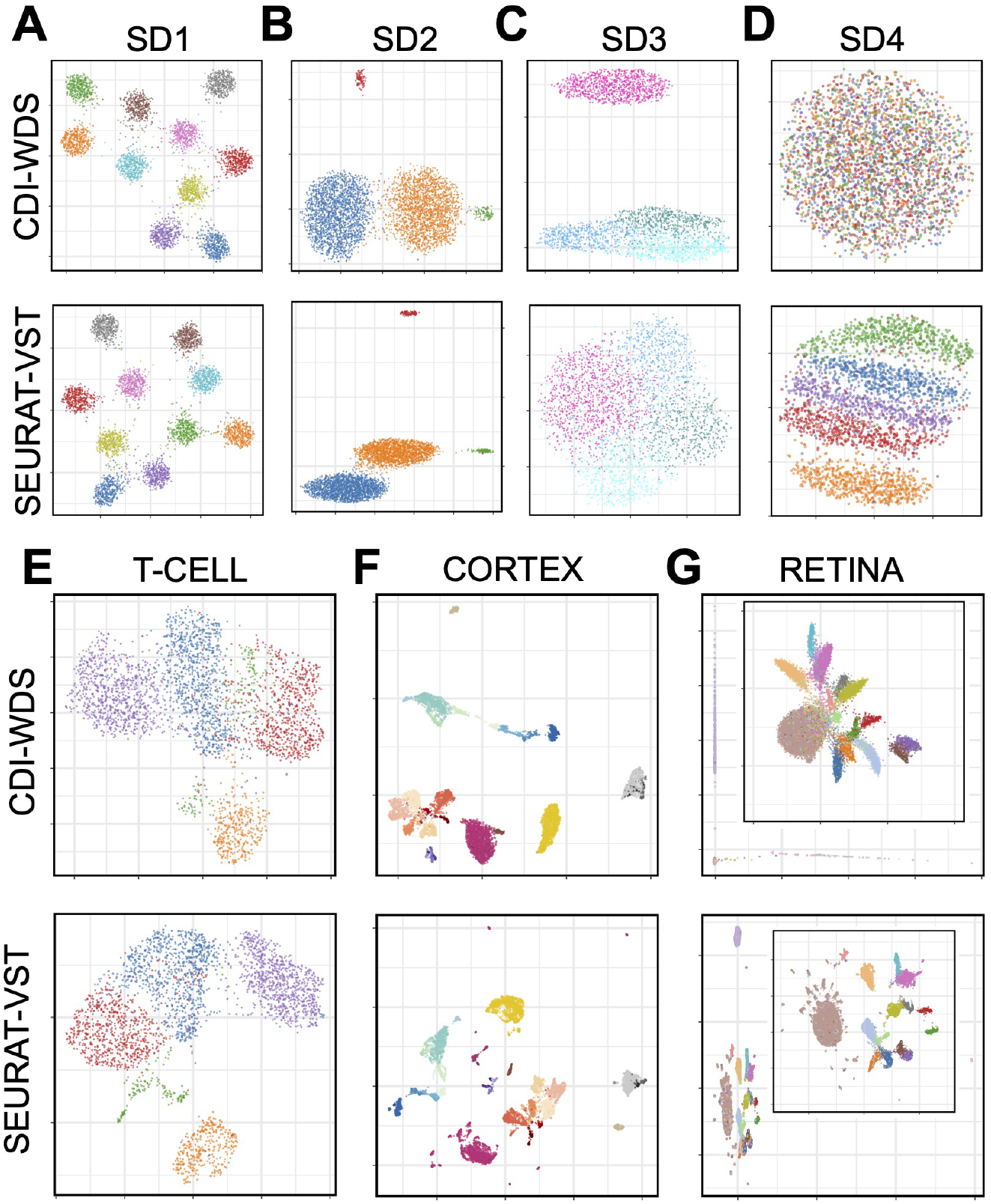
Comparisons between WDS and VST. (*A*)–(*D*) UMAPs of the simulated datasets (SD1–SD4); (*E*)–(*G*) UMAPs of the experimental datasets (T-CELL, CORTEX, and RETINA). In all plots, cells from different cell types are marked with different colors.

- For all datasets except for SD3 and SD4, the UMAPs based on both the WDS-selected feature genes and the VST-selected feature genes separate different cell clusters well.
- For SD3, WDS selected 74/75 of the actual feature genes while VST only selected 48/75. Most of the feature genes missed by VST but not WDS are highly expressed in three similar subtypes but lowly expressed in the less similar main types. Consequently, the UMAP based on the WDS-selected feature genes reflects the cell type structure better than the VST-selected feature genes.
- For SD4, VST performed better than WDS. SD4 was generated from Splatter (Zappia et al. 2017), a scRNA-seq data simulator that imposes strong mean-dispersion trends on gene expressions – that is, highly expressed genes are forced to have a lower dispersion. Such trends are commonly seen in bulk RNA-seq data, but we did not observe them in the UMI counts of the scRNA-seq datasets (Supplemental Fig. S9). On datasets with such trends, WDS will select genes with lower average UMI counts. These genes contain little information on cell types; thus, the resulting UMAP cannot separate the cells from different cell types. Because splatter is a commonly used scRNA-seq data simulator, we included SD4 to check the robustness of the subsequent procedure of CDI. In practice, when such mean-dispersion trends exist for the UMI counts, we should not use WDS to select feature genes; however, when such mean-dispersion trends do not exist (as in all of our experimental datasets), WDS works well.

When WDS and VST select different feature gene sets, even if their resulting UMAPs separate cell types similarly well, CDI based on the two gene sets could select different label sets. For example, for T-CELL, both UMAPs look similar (Fig. 3E). However, CDI following VST selected the six-cluster label set generated by the spectral clustering (ARI=0.39); CDI following WDS selected the five-cluster label set generated by Seurat (ARI=0.87). For reference, T-CELL’s benchmark label set has five clusters, similar to the five-cluster label selected by CDI following WDS (T-CELL panel in Fig. 4A, B). Thus, CDI following WDS is more robust and accurate.

**Figure 4.**
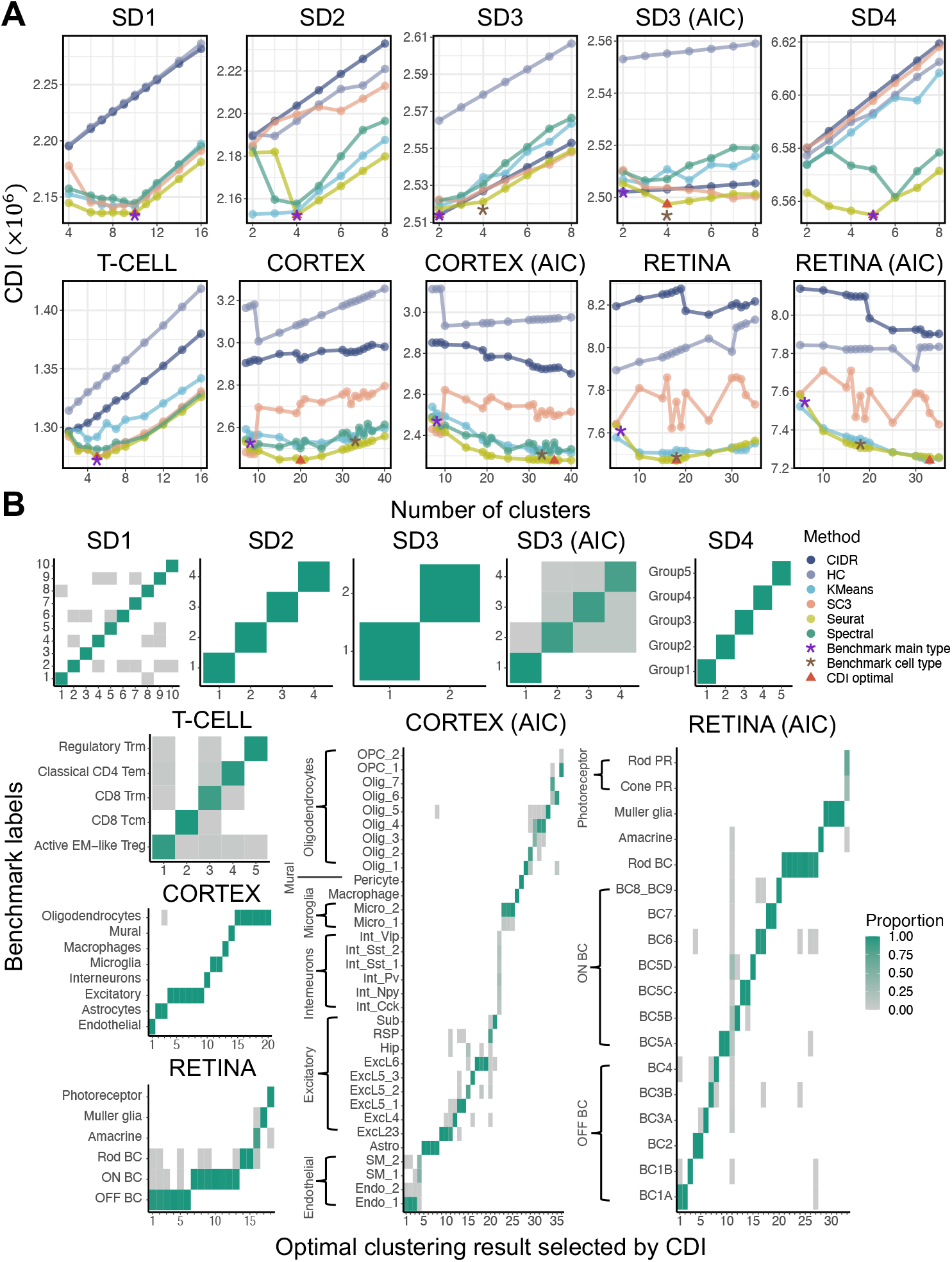
CDI’s performance evaluation. CDI was applied to four simulated datasets (SD1-SD4) and three experimental datasets (T-CELL, CORTEX, RETINA). All datasets have benchmark label sets. If the subplot title contains AIC, CDI-AIC was applied; otherwise, CDI-BIC was applied. (*A*) CDIs of different candidate label sets. The x-axis labels the cluster number in the candidate label sets, and the y-axis labels their corresponding CDIs. For datasets with hierarchical structures (SD3, CORTEX, and RETINA), both CDI-BIC and CDI-AIC were applied. The line color represents the clustering method. The red triangle marks the CDI-selected label set with the corresponding clustering method, the number of clusters, and the CDI value. The purple star marks the benchmark main type label set. The brown star marks the benchmark subtype label set (if available). (*B*) Benchmark cell proportion heatmaps of different candidate label sets. The x-axis labels the clusters in the candidate labels set, and the y-axis labels the benchmark cell types. The color of each rectangle represents the benchmark cell type proportions in each candidate cluster. Each column adds up to 1. PR, photoreceptor.

When WDS selects different numbers of feature genes, CDI based on these different feature gene sets has robust performance. For example, for T-CELL, based on the 200 WDS-selected feature genes, CDI selected a six-cluster label set generated by Seurat; the second-best was the five-cluster label set generated by Seurat. Based on the 300 WDS-selected feature genes, CDI selected the five-cluster label set generated by Seurat. Finally, based on the 400 or 500 WDS-selected feature genes, CDI selected the five-cluster label set generated by SC3. (Supplemental Fig. S14). These label sets were all similar to the benchmark label set (ARIs between 0.80 and 0.87). Thus, CDI’s performance is robust to the number of WDS-selected feature genes.

#### 2.4.2 Performance of CDI in selecting the optimal label set

We evaluated the performance of CDI using the candidate label sets generated by multiple clustering methods, each with a wide range of tuning parameters. The clustering methods we applied include hierarchical clustering, K-means clustering, spectral clustering, CIDR (Lin et al. 2017), Seurat V3 (Stuart et al. 2019), and an ensemble clustering method called SC3 (Kiselev et al. 2017).

##### A. Data containing no rare cell types

We define a cell type as *rare* if its proportion is below 3%. We evaluated the performance of CDI on the datasets where none of the cell types are rare. These datasets include SD1, SD4, and T-CELL. SD1 contains ten equally proportional cell types simulated from the NB model verified in Section 2.2; SD4 contains five unequally proportional cell types simulated from Splatter; T-CELL contains a mixture of five cells types of T cells. We selected their feature genes with either WDS (SD1, T-CELL) or VST (SD4) and then applied CDI-BIC to select the optimal label set marking their main cell types. CDI-BIC performed very well on all three datasets. It selected the label sets with the correct numbers of clusters (Fig. 4A); moreover, the selected label sets are very similar to the benchmark label sets (Fig. 4B). Of note, some other candidate label sets have the correct numbers of clusters, but their cell labels are very different from the benchmarks. For example, for T-CELL, two label sets with the lowest CDI are the five-cluster label sets generated by Seurat and SC3. These two label sets have similarly low CDIs (1.2744 × 10^6^ for Seurat and 1.2743 × 10^6^ for SC3) and similarly high ARIs (0.875 for Seurat and 0.870 for SC3). However, the five-cluster label sets generated by other methods have much higher CDIs (1.3356×10^6^, 1.3095×10^6^, 1.2918×10^6^, 1.2812×10^6^) and much lower ARIs (0.002, 0.116, 0.132, and 0.388). Apparently, the label sets with lower CDIs have higher ARIs. The heatmaps to compare the candidate label sets with the benchmark label sets also verified that the label sets with lower CDIs are more similar to the benchmark label set (Supplemental Fig. S10). These results suggest that CDI has similar performance with ARI in selecting the optimal label set when the data contain no rare cell types. Moreover, CDI has a significant advantage over ARI because its calculation does not rely on the knowledge of the benchmark label set.

##### B. Data containing rare cell types

We evaluated the ability of CDI-AIC and CDI-BIC to detect rare cell types. For example, SD2 simulated a cell population with two normal-sized (47.62% of all cells each) and two rare (2.38% of all cells each) cell types. Two normal-sized cell types and one rare cell types have different but similar feature gene UMI count distributions. This rare cell type is called RC1; the other rare cell type is called RC2 (Fig. 3A SD2 panel). For SD2, CDI-BIC selected a label set similar to the benchmark, suggesting that it can differentiate both rare cell types. Next, we reduced the cell proportion of RC1 further to challenge CDI with more difficult tasks. When the cell proportion of RC1 reduced to 2.03% (85/4185), CDI-BIC selected the three-cluster label set generated by spectral clustering; however, CDI-AIC still selected the four-cluster label set including RC1 (Supplemental Fig. S13B, S13D). When the cell proportion of RC1 reduced to 0.49% (20/4120), neither CDI-AIC nor CDI-BIC distinguished RC1; instead, they both selected the three-cluster label set generated by Seurat. Although this label set misses RC1, it has a very high ARI (0.98) (Supplemental Fig. S13G, S13I). Also, if we put the benchmark label set to the candidate pool, CDI-BIC would rank it as the fourth among all label sets, and CDI-AIC would select it as the optimal (Supplemental Fig. S13I). These results suggest that both CDI-BIC and CDI-AIC perform well in detecting rare cell types; compared with CDI-BIC, CDI-AIC is more sensitive in detecting rare cell types.

##### C. Data with hierarchical cell type structures

In scRNA-seq data, some main cell types can be further divided into subtypes. For example, SD3 simulated a cell population with two main cell types; one main cell type contains three subtypes, and the other is homogeneous. We used CDI-BIC to select the main type label set and CDI-AIC to select the subtype label set. As a result, CDI-BIC selected the two-cluster label set similar to the benchmark main type label set; CDI-AIC selected the four-cluster label set similar to the benchmark subtype label set (Fig. 4A SD3 panel, 4B SD3 panel). Another dataset, CORTEX, has eight main types and 33 subtypes. CDI-BIC selected a 20-cluster label set (Fig. 4A CORTEX panel). These 20 clusters correspond to the partitions of eight benchmark main types: clusters 4-10 correspond to excitatory neurons, clusters 11 and 12 correspond to microglia cells, and clusters 15-20 correspond to oligodendrocytes (Fig. 4B CORTEX panel). CDI-AIC selected a label set with 36 clusters (Fig. 4A CORTEX (AIC) panel). Some of these 36 clusters were partitions of the benchmark subtypes: they further partitioned Endothelial cells subtype 1, Astrocytes, Excitatory cells subtype 23, Excitatory neuron 5_1, Excitatory neuron 6, Oligodendrocytes subtype 5, and microglia subtype 2 into 21 clusters. Other clusters in the 36-cluster label set were mixtures of rare cell types, including a cluster mixing all interneuron subtypes (taking up 1.8% of all cells), a cluster mixing two Endothelial subtypes, and a cluster mixing two Microglia subtypes (Fig. 4B CORTEX (AIC) panel) In fact, not only the selected label set but also all other candidate label sets cannot separate these rare cell types. These results suggest that when the data have hierarchical structures, applying CDI-BIC in combination with CDI-AIC is an excellent strategy to reveal its hierarchical structure. When the subtypes contain too few cells, CDI-AIC may fail to identify the rare subtype but can still cluster them with other similar subtypes.

##### D. Data from multiple batches

RETINA had two batches with six main types. Among them, photoreceptors were further divided into rod photoreceptors (0.34%) and core photoreceptors (0.18%. The ON cone bipolar cells (BCs) had seven subtypes; the OFF cone BCs had six. CDI-BIC selected an 18-cluster label set that classified the main types well. It further partitioned the rob BCs, ON cone BCs, and OFF BCs into several subtypes. On the other hand, CDI-AIC selected a label set with 33 clusters. This label set separated all the subtypes well. Besides, it provided a more exemplary partition on the benchmark Müller glia cells and the rod bipolar cells (RBCs). Some subtypes of ON cone BCs (BC5A, BC5C, BC6, BC7) and OFF cone BCs (BC1A, BC2) were also further partitioned into sub-clusters. Cluster 11 spanned many cell types; however, they are mainly BCs. These results suggest that CDI works well on the multi-batch scRNA-seq data by incorporating hypothesis tests for significant batch effect and subsequent adaptive modeling (Section 4.1.2).

### 2.4.3 Comparison of CDI with other unsupervised indices

For general clustering problems, many other unsupervised indices are developed, including Davies-Bouldin index (Davies and Bouldin 1979), Silhouette coefficient (Rousseeuw 1987), and Calinski-Harabasz index (Caliński and Harabasz 1974). Although these methods are not customized for scRNA-seq data clustering, they have been applied in multiple publications to select clustering label sets (H Jiang et al. 2018; Peyvandipour et al. 2020; Liu et al. 2020). We compared the performance of these unsupervised indices and CDI by using the benchmark supervised index ARI. An unsupervised index performs well if it is a monotone function of ARI.

Because CDI-BIC aims to select main type label sets, we compared it with the ARI using the benchmark main type label sets. Similarly, we compared CDI-AIC with the ARI using the benchmark subtype label sets. Both CDI-AIC and CDI-BIC show clear monotone trends with ARI in all the datasets (Fig. 5). Therefore, CDI has matching performance with ARI, which requires the prerequisite knowledge of the benchmark label sets.

**Figure 5.**
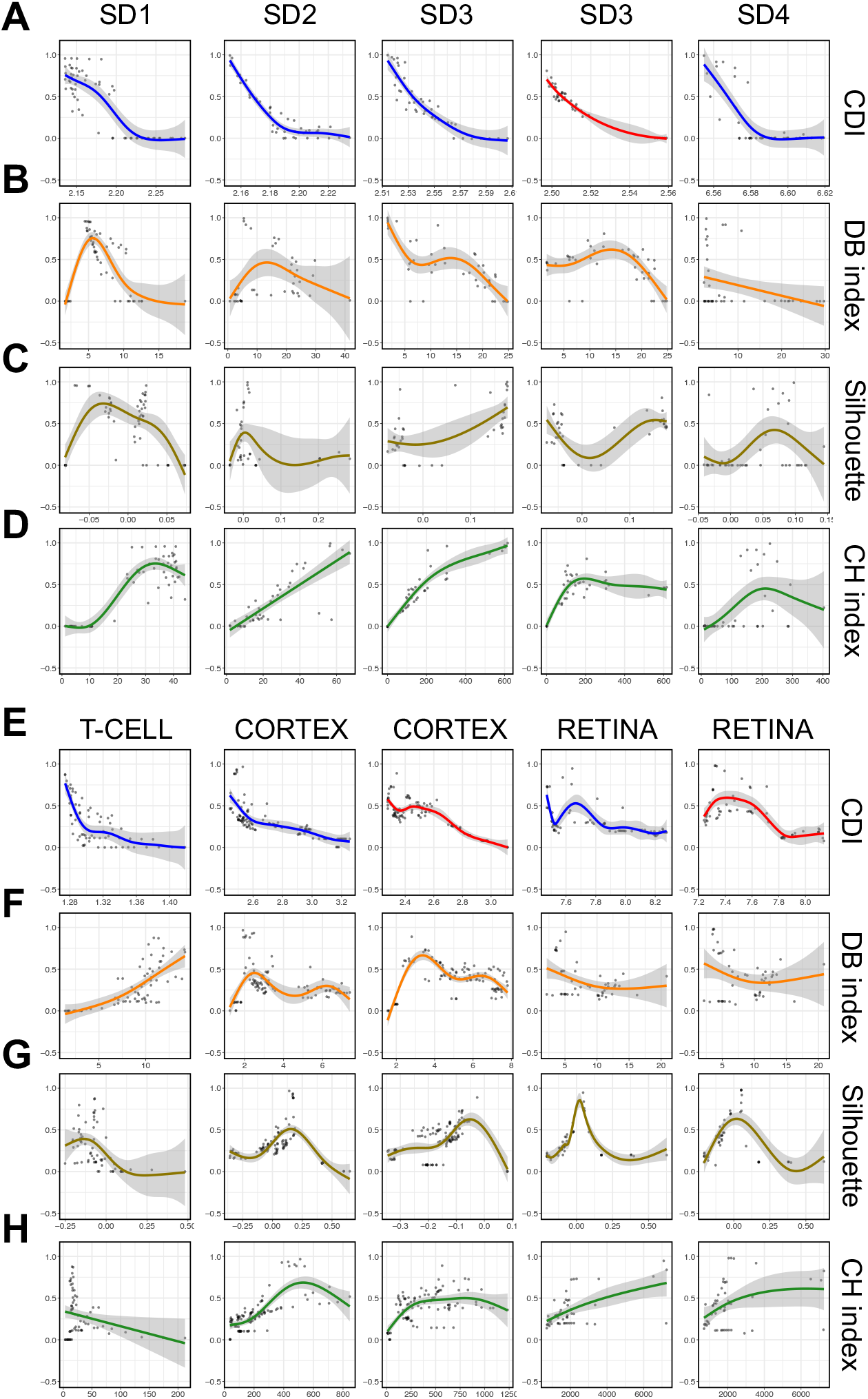
Comparison between CDI and other unsupervised indices. In each panel, the x-axis is an unsupervised index; the y-axis is ARI; each dot represents a candidate label set as introduced in Fig. 4A. (*A*)–(*D*) are for simulated datasets, and (*E*)–(*F*) are for experimental datasets. (*A*) and (*E*) ARI vs. CDI (blue for CDI-BIC and red for CDI-AIC). (*B*) and (*F*) ARI vs. Davies-Bouldin index. (*C*) and (*G*) ARI vs. Silhouette coefficient. (*D*)and (*H*)ARI vs. Calinski-Harabasz index. The curves are fitted using these dots based on the generalized additive model (Hastie 2017), and their 95% confidence bands are shown using the grey regions around the curves.

In contrast, other unsupervised indices did not exhibit any apparent monotone relationships with ARI. They sometimes assigned similar scores to two label sets with very different ARIs. For example, the Davies-Bouldin index assigned similar scores to many varying-quality label sets in SD1 and CORTEX (Fig. 5B, F). Same for the Silhouette coefficient in SD1, SD2, and T-CELL (Fig. 5C, G) and the Calinski-Harabaz index in T-CELL (Fig. 5H). Therefore, these general unsupervised indices are inferior to CDI in selecting the optimal clustering label set for scRNA-seq data.

## 3 Discussion

In this study, we develop a new index, CDI, to calculate the deviation between the candidate label set and the observed UMI counts. For each candidate label set, CDI calculates the negative penalized maximum log-likelihood of the feature gene UMI counts. The likelihood function is calculated based on the gene-specific cell-type-specific NB distribution family verified in both monoclonal and polyclonal scRNA-seq data. We recommend using WDS to select the feature genes for CDI because CDI following WDS has robust and satisfying performances.

Because calculating CDI relies on the gene-specific cell-type-specific NB distribution family, we would like to elaborate on two major innovations of our approach to ensure the distribution family is reliable.

First, to generate a monoclonal dataset, we cloned a single mother cell to derive a cell line whose components can be considered identical. This strategy is better than the existing artificial External RNA Control Consortium (ERCC) or fluorescence-activated cell sorting (FACS) strategies for the following reason. For scRNA-seq data, variations in UMI counts could come from the variations in the efficiency of library construction, sequencing depths, cell cycles, and cell types. The first three variations cannot be avoided even in a monoclonal scRNA-seq dataset, while the last one is successfully eliminated. Monoclonal single cell datasets are essential to characterize the UMI count distributions in scRNA-seq data: with the well characterized distributions, we can use the model validation tools such as AIC or BIC to evaluate the deviation from the data to the model given the candidate label set. Previously, such monoclonal datasets were generated by either piking in the ERCC RNA or purifying cells by FACS. The ERCC RNA samples contain the synthesized RNAs differing from the endogenous transcripts in many aspects (Zheng et al. 2017; L Jiang et al. 2011) (such as length, guanine-cytosine content, 5’ cap, polyA length, and ribosome binding). These structural disparities lead to different conversion efficiencies of mRNA into cDNA. Thus, while ERCC eliminates the cell type variations, they also eliminate or distort the variations in library construction, sequencing depths, and cell cycles. Another choice, FACS, can keep these variations; however, it only purifies cells based on a limited number of protein markers and therefore can only reduce but not eliminate the cell type heterogeneity in a cell population. Different from these two existing strategies, we used the single-cell expansion strategy to ensure an ideal monoclonal population: it keeps the variations in library construction, sequencing depths, and cell cycles; on the other hand, it also eliminates cell type heterogeneity.

Second, although we are not among the first to suggest that the UMI count distributions are not necessarily zero-inflated, we characterized the UMI count distributions with more specific gene-specific and cell-type-specific models. Previous studies model the UMI count distributions differently (Townes et al. 2019; Svensson 2020). Townes et al. (2019) models the distributions of cellular gene UMI counts as an over-dispersed Dirichlet-multinomial distribution, which can be approximated by the independent NB models with gene-common dispersion parameters. Svensson (2020) proposes to use the NB models with gene-common and cell-type-common dispersion parameters. Neither evaluated or compared the proposed model with other candidate NB models. To derive a reliable NB distribution family, we used both monoclonal and polyclonal datasets to evaluate the fitting of distribution families to the UMI count distributions. Eventually, we found that the NB distribution family with the gene-specific cell-type-specific mean and dispersion parameters is the best.

Next, we would like to elaborate on the key features and limitations of CDI.

First, calculating CDI relies on the likelihood of the raw UMI counts. If only the normalized UMI counts are available, CDI cannot be applied.

Second, CDI is not a clustering method; instead, it is an index to evaluate the quality of the candidate label set: the label set with the lowest CDI will be selected. Thus, the quality of the selected label set highly depends on the quality of the candidate label sets. If none of the candidate labels fits the data well, the CDI-selected label set will not improve the fitting. Thus, providing a large pool of candidate label sets is crucial. We suggest using at least three methods, each with at least five labels with different cluster numbers.

Third, CDI-AIC puts fewer penalties on cluster numbers and usually selects the label with more clusters than CDI-BIC. These clusters often correspond to cell subtypes, and some could mark the cell transition stages. In many cases, the collected cells have different developmental stages so that the scRNA-seq data exhibit trajectory patterns. In those datasets, we can consider the cell types as “continuous”. When discrete clustering methods are applied to these datasets, the resulting clusters often represent a local stage on the trajectory. Although CDI is not designed to select the optimal trajectory, we can use it to select the label sets corresponding to the optimal local stages. For example, in RETINA, the subtypes of ON cone BCs and OFF cone BCs represent different development stages of those BCs. The CDI-AIC successfully selected a satisfying discrete label set that approximates the continuous trajectories of ON cone BCs and OFF cone BCs (Fig, 4B. RETINA (AIC)).

In summary, finding an optimal clustering label set for scRNA-seq data is critical because clustering impacts all downstream analyses; CDI provides a robust and accurate unsupervised method to select cluster label sets and hence contributes to the reliability of downstream scRNA-seq analysis.

## 4 Methods

We develop CDI in two stages. In stage I, we characterize the unique molecular identifiers (UMIs) count distributions using the experimentally generated monoclonal and polyclonal cell populations. In stage II, based on the UMI count distributions in stage I, we develop the CDI and select the label set with the lowest value as the optimal label set.

### 4.1 Models to characterize the UMI counts

#### 4.1.1 Single-batch UMI count distributions

In a single batch scRNA-seq dataset, denote the UMI count for gene *g* and cell *c* by *X_gc_, g* ∈ {1,…, *G*}, and *c* ∈ {1,…, *N*}. Suppose the *true* label of the *N* cells is ***L***_0_ = (*L*_0,1_, *L*_0,2_,…, *L*_0,*N*_)′ with *L*_0,*c*_ ∈ {1, 2,…, *K*_0_}. Here *K*_0_ is the number of the underlying true cell types.

Based on our experiments and modeling, assume

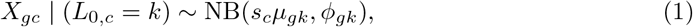

where *s_c_* is a scale factor to adjust for the imbalance of the cellular total UMI counts in the scRNA-seq data (Supplementary Note 1), and *μ_gk_* and *ϕ_gk_* are the mean and dispersion parameters of gene *g* in cell type *k*. The probability mass function of negative binomial distribution is shown in Supplementary Note 1.

The cellular scale factor *s_c_* only impacts the mean but not the dispersion of the NB distribution. From a Bayesian’s perspective, we can view *X_gc_* as a Poisson-gamma distribution where

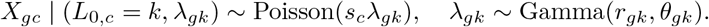

Then, (1) holds with *μ_gk_* = *r_gk_θ_gk_* and *ϕ_gk_* = 1/*r_gk_*.

#### 4.1.2 Multi-batch UMI count distribution

The UMI count distributions may differ across batches. When multiple batches exist, we model the UMI count distribution in batch *b* as

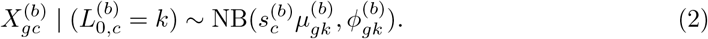

We assume batch effect lies in a low-dimensional space. Thus, for some genes or some cell types, introducing batch-specific parameters might not be necessary.

To decide whether to introduce batch-specific parameters for a gene in a cell type, we first test the hypothesis:

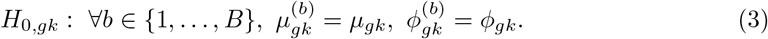

We used the likelihood ratio test to test this hypothesis. If the hypothesis is rejected, it indicates that the batch effect is significant for the gene in this cell type; thus we will introduce the batch-specific parameters like in (2) to model the UMI count of this gene in this cell type. Otherwise, even if the batch effect exists, it is not significant; thus we will not use the batch-common model (1) to characterize the UMI count distribution of this gene in this cell type.

### 4.2 Cell-type-specific “goodness-of-fit” tests

We filtered the genes and the cells as described in Section 4.6. We derived a “goodness-of-fit” test for each gene. First, we assigned the cells into 5*K*_0_ bins, where 5 is the number of UMI count categories and *K*_0_ is the number of the benchmark cell type. Based on the values of (*X_gc_, L_c_*), we assigned cell c into one of the following bins:

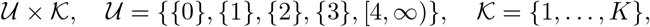

where × is the Catesian product of two sets. Second, we computed the test statistic as

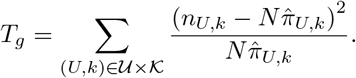

Here, 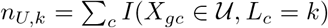, and 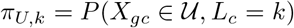, which are the parameters of the multinomial distributions on 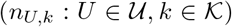. Because *π_U,k_* is unknown, we estimated π_U,k_ by first expressing it as a function of (*μ_gk_, ϕ_gk_*) in the corresponding NB distribution family and then derived the maximum likelihood estimator (MLE) in the multinomial likelihood (Chernoff and Lehmann 1954). Third, if *T_g_* is larger than the 95% quantile of the chi-square distribution with the degree of freedom 5*K*_0_ – *p* – 1, we rejected the “goodness-of-fit” hypothesis for gene *g*. Here, *p* is the number of parameters in the corresponding NB distributions: for cell-type-common NB distributions, *p* = 2; for cell-type-specific NB distributions, *p* = 2*K*_0_. We used the chi-square quantile as the threshold because when the UMI count of gene *g* follows the corresponding NB model and *L_c_* all match the true cell types, *T_g_* asymptotically follows *χ*^2^(5*K*_0_ – *p* – 1) (Chernoff and Lehmann 1954). Finally, we performed the test for all the genes and calculated the rejection proportion. We used the proportion as the criterion to assess the overall fitting of the corresponding NB distribution family and the UMI count distribution.

### 4.3 Working dispersion score (WDS) to select feature genes

WDS is a score to measure the average sample (cell) dispersion of each gene.

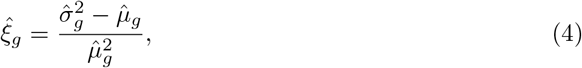

where 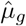 and 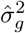 are the sample mean and variance of gene *g*’s UMI counts in the pooled data. It can be viewed as the estimator of 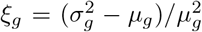, where *μ_g_* and 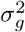 are the mean and variance of gene *g*’ UMI counts. The rationale of WDS is provided in the supplementary note 2.

Before selecting the feature genes, we filtered the genes with more than 95% zero counts. We computed WDS for each gene and selected the top *G*_1_ (default *G*_1_ = 500) genes with the highest WDS. For datasets with multiple batches, we calculated and ranked WDS for genes in each batch: the minimum rank across the batches is set as the overall rank for each gene. In the end, we selected the top *G*_1_ (default *G*_1_ = 500) genes with the highest overall rank as the feature genes.

In our software, feature genes can also be added manually. For example, the users can use the feature genes provided by other approaches or domain knowledge.

### 4.4 CDI optimal label selection

Denote a label set with *K* cell types by ***L*** = (*L*_1_,…, *L_N_*)′ with *L_c_* ∈ {1,…, *K*}. This label set is not necessarily the true label set. It could be derived from any clustering method.

Let 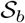 be the set of cells that belong to batch *b*. Let 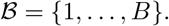, and 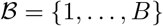.

Based on model 2, if label sets are accurate, then the likelihood function of all genes and all cells are

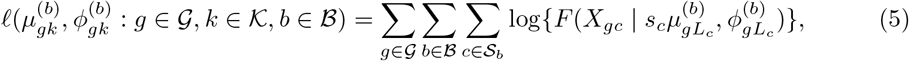

where *F* is the probability mass function of negative binomial distribution (Supplementary Note 1). Across *B* batches, for each feature gene *g* and cell type *k*, we use the score tests to check if batch-specific mean and dispersion parameters need to be introduced. If *H*_0,*gk*_ is accepted, we will use a batch-common but cell-type-specific mean and dispersion parameters for this feature gene, *i.e*.,

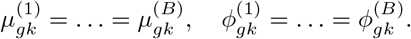

If *H*_0,*gk*_ is rejected, we will introduce batch specific and cell-type-specific mean and dispersion parameters.

Next, we obtain the MLE 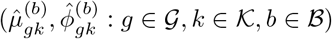 based on (5). The corresponding 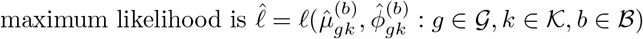.

To adjust for the model complexity, we use the penalized negative log-likelihood function as CDI

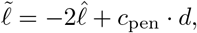

where *c*_pen_ is the scale of penalty, and *d* is the overall degree of freedom of the model. For AIC, *c*_pen_ = 2; for BIC, *c*_pen_ = log(*N*). The overall degree of freedom 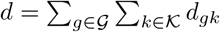. If *H*_0,*gk*_ is rejected, *d_gk_* = 2*B*; otherwise, *d_gk_* = 2.

In practice, when *T* label sets {***L**_t_*: *t* ∈ {1,…, *T*}} are given, we calculate the CDI 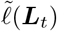 for each label set ***L**_t_*. If ***L**_t_* is accurate, 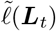 tends to be small. The optimal label set is chosen by 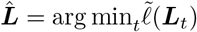.

Because BIC adds more penalty on the model degree of freedom, it prefers the models with fewer numbers of clusters. Thus, we use CDI-BIC to select the main type label set and CDI-AIC to select the subtype label set.

### 4.5 Simulation setting

We simulated three sets of single-cell data (SD1-SD3) from the Negative Binomial distribution with gene-specific and cell-type-specific parameters. More specifically for SD1-SD3, the gene expression level for cells in cell type *k* and gene *g* were randomly sampled from NB(*μ_gk_, ϕ_gk_*), where *μ_gk_* represented the mean parameter and *ϕ_gk_* represented the dispersion parameter. Each dataset contained 10,000 genes, and the number of cells ranged from 2,800 to 4,200. To test the robustness of CDI, we generated SD4, which contained many outliers, and the UMI count distributions no longer followed the verified NB distribution.

**SD1.** We generated ten equal-sized cell groups. Each group contained 400 cells; thus, in total there are 4,000 cells. One cell type was treated as the baseline type with *μ_gk_* generated from the truncated normal distribution with mean 0.2 and standard deviation 0. 1, and *ϕ_gk_* generated from the truncated normal distribution with mean 0.5 and standard deviation 0.1. For each of the other nine groups, 25 genes had mean parameters shifted from the baseline group with log2 fold change 2.4. The feature gene dispersion parameters were shifted by a Gaussian-distributed factor with mean 0 and standard deviation 0.05.
**SD2.** We generated two normal-sized cell types with 2,000 cells each and two rare cell types with 100 cells each. One normal-sized cell type was treated as the baseline group with *μ_gk_* generated from the truncated normal distribution with mean 0. 2 and standard deviation 0.1. The other normal-sized cell type contained 40 feature genes with log2 fold change of mean 1.5. One rare cell type, RC1, contained 50 feature genes with log2 fold change of mean 2.8; the other rare cell type, RC2, contained 50 feature genes with log2 fold change of mean 3.2. Because the log2 fold change of RC1 is smaller than RC2, RC1 is considered to be more similar to the two normal-sized cell types. The dispersion parameters *ϕ_gk_* were set in the same way as in SD1.
**SD3.** We generated two main-types: C1 contains 1, 000 cells from a homogeneous cell type, and C2 contains 1, 800 cells from 3 subtypes. C1 is the baseline group with *μ_gk_* generated from the truncated normal distribution with mean 0.4 and standard deviation 0.1 and *ϕ_gk_* generated from the truncated normal distribution with mean 1 and standard deviation 0.1. Each subtype of C2 contains 600 cells and 40 feature genes. Among these 40 feature genes, 30 were shared by all subtypes, and the rest 10 were exclusive for each subtype. The log2 fold change in the means was 4 for the main type and 1.8 for the subtype. The dispersion parameters *ϕ_gk_* were set in the same way as in SD1.
**SD4.** We generated five common cell types using the R package Splatter (Zappia et al. 2017) with 3, 000 cells and 5, 000 genes. The probabilities that a cell belongs to any cell group were 0. 2 for all groups. The proportion of differentially expressed genes were 1% per cell group. The location and scale parameters of the log-normal distribution for these feature genes were (0.4, 0.1). In addition, we followed the default option to add 5% of outliers. After we filtered the cells with less than 1% non-zero counts and the genes with less than 1% of non-zero cells, the dataset had 4, 887 remaining genes. See Supplemental Fig. S12 for all the parameters used in this setting.

### 4.6 Description of the experimental scRNA-seq datasets and their preprocessing

**CT26.WT.** This dataset was generated in Dr. Qi-Jing Li’s lab. Dr. Li is one of the co-authors of this paper. The wild-type CT26 cells from the murine colorectal carcinoma cell line were single-cell-diluted, and a clone was picked and cultured for 220 days. For the single-cell RNA-seq library preparation, 10, 000 cells of each clone were processed with the protocol of Chromium Single Cell 3’ Reagent kits v3 from 10X Genomics to make the single-cell RNA sequence library. Cells with more than 10% mitochondrially derived transcripts were removed. Among these cells, we selected those with non-zero gene proportions greater than 3% or the number of non-zero genes greater than 300 (at least one of the two conditions needed to hold). We further selected genes with non-zero count proportions greater than 1% or the number of non-zero cells greater than 50. This dataset was of high quality, with 24, 208 median UMI counts per cell and 4, 376 median genes per cell. Since this dataset was highly homogenous, we used CT26.WT to evaluate the Pearson’s chi-squared “goodness-of-fit” of different models to the UMI counts in the monoclonal scRNA-seq data.
**T-CELL.** The T-CELL dataset was generated in our previous study (Christian et al. 2021). The benchmark clustering labels of the T-CELL population were generated as a combination of protein-marker-based flow sorting labels and bioinformatics labels from Seurat v2. For evaluation purpose, we selected 5 distinct cell types: Regulatory Trm cells, Classical CD4 Tem cells, CD8 Trm cells, CD8 Tcm cells, and Active EM-like Treg cells. In this study, tumors were firstly collected from the female mice after 3 weeks since the mice were injected by 4T1 tumors. Tissues were then disassociated into single cells and homogenized. T cells were separated out by flow sorting with a stringent gating threshold and sequenced on the 10X platform. For preprocessing, we filtered out genes with less than 2% non-zero cells and removed cells with less than 2% non-zero genes. Eventually, 2, 989 cells from five cell types with 7, 893 genes were retained.
**CORTEX.** The visual cortex dataset was generated by Hrvatin et al. (Hrvatin et al. 2018) using inDrop to study the diversity of activity-dependent responses across cortical cell types. We obtained the labeled scRNA-seq dataset from Huang et al. 2018, which contained 10, 000 cells with 19, 155 genes. Among these 10, 000 cells, 7, 390 cells were identified to 33 cell types as an intersection of Seurat v1 and a density-based method (Rodriguez and Laio 2014). In addition, eight main cell types (excitatory neurons, oligodendrocytes, astrocytes, interneurons, etc.) were annotated with known feature genes. We selected cells with at least 300 or 3% of non-zero genes, and genes with at least 50 or 1% non-zero cells. After preprocessing, 7, 376 cells with 12, 887 genes were included in clustering.
**RETINA.** The mouse retina dataset was generated by Shekhar et al. 2016 using Drop-seq to classify retinal bipolar neurons. This dataset contained 27,499 cells with 13,166 genes. Among these 27, 499 cells, 26, 830 cells were labeled with 18 cell types by the assembled pipeline: they first used Louvain-Jaccard (Blondel et al. 2008) method to cluster the cells and then annotated the clusters with known feature genes. This 18-cluster label set was treated as the benchmark subtype label set. We further grouped the cell types into 6 main types based on the original paper. These cells came from two experimental batches of FAC sorted Vsx2-GFP positive cells on different days. We selected cells with at least 300 or 3% of non-zero genes, and genes with at least 100 or 2% non-zero cells. After preprocessing, all 26, 830 cells with 13,118 genes were selected. The preprocessing step removed very few genes and cells because the dataset obtained from the original paper was filtered before cell type annotation.

## 5 Data access

ScRNA-seq for the murine colorectal carcinoma cell line CT26.WT is available at https://data.mendeley.com/datasets/cb2cb8f8mp/2. Three other public scRNA-seq datasets were also used in this study: T-CELL (https://data.mendeley.com/datasets/3f4rsk96kf/3), CORTEX (GSE102827), and RETINA (GSE81905).

We used the CDI R packager version 0.99.2 in this study. This R package has been submitted to the Bioconductor project. The development version of CDI is available from https://github.com/jichunxie/CDI. The scripts for to reproduce figures of the manuscript using this package are available at https://github.com/jfanglovestats/CDI_figures.

## 6 Competing interest statement

The authors declare no competing interests.

## 7 Acknowledgements

Ms. Jiyuan Fang’s research was partially supported by Duke Center for Human System Immunology (CHSI). Dr. Cliburn Chan’s research was partially supported by the Duke University Center for AIDS Research (NIH fund 5P30 AI064518) and the Duke Senescent Cell Evaluations in Normal Tissues (SCENT) Mapping Center (NIH fund 1U54AG075936-01). Dr. Kouros Owzar and Dr. Liuyang’s research was supported by Duke University. Dr. Diyuan Qin’s research was supposed by Duke University. Dr. Qi-Jing Li’s research was partially supported by NIH Award R33 CA225328. Dr. Jichun Xie’s research was partially supported by SCENT (NIH fund 1U54AG075936-01). In addition, Drs. Cliburn Chan, Qi-Jing Li, and Jichun Xie’s research was partially supported by the Translating Duke Health (TDH) Controlling the Immune System grant. We also thank the Duke CHSI for providing the computation resources and administrative support.

The content is solely the responsibility of the authors and does not necessarily represent the official views of the National Institutes of Health.

## Notes

https://data.mendeley.com/datasets/cb2cb8f8mp/2

### Competing Interest Statement

The authors have declared no competing interest.

