## Supplemental materials for "Clustering Deviation Index (CDI): A robust and accurate unsupervised measure for evaluating scRNA-seq data clustering"

### 1 Supplementary Figures

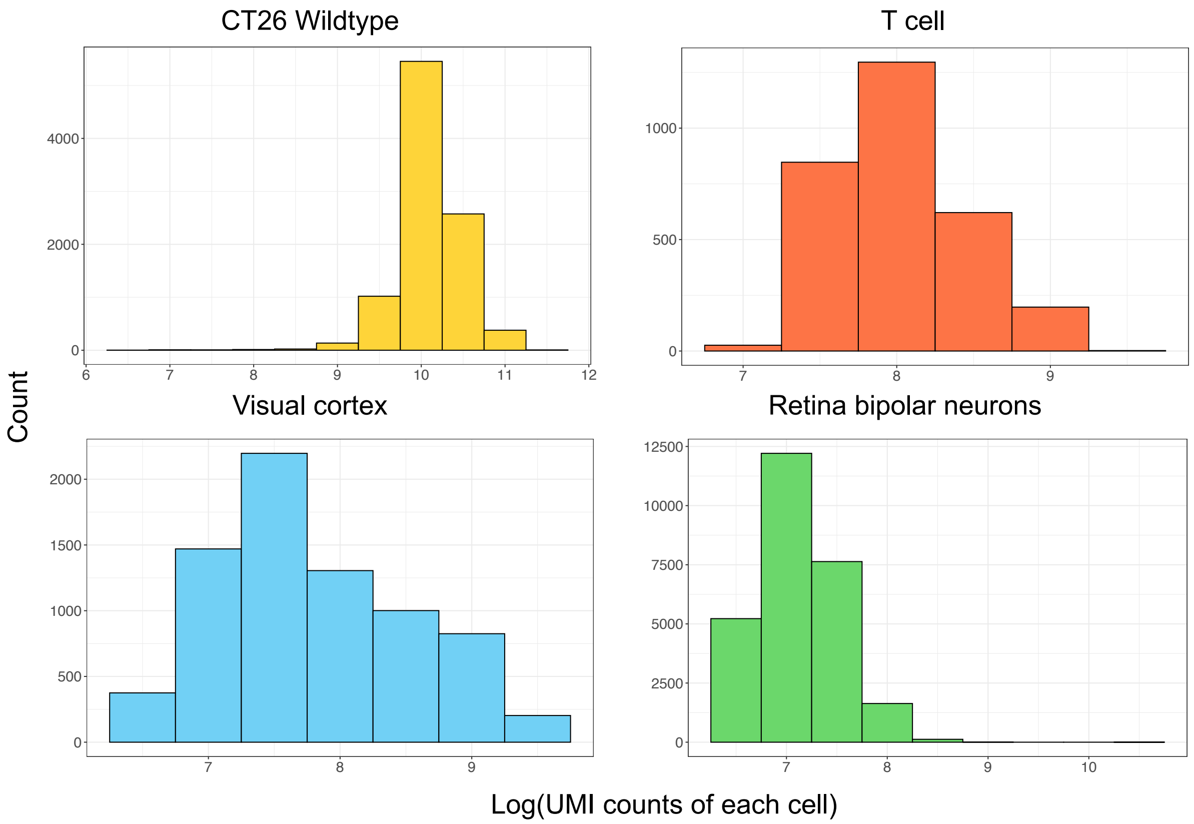


**Supplementary Figure 1: Histogram of UMI counts.** The x-axis is the log-transformation of total UMI counts for each cell. The y-axis is the count of cells.

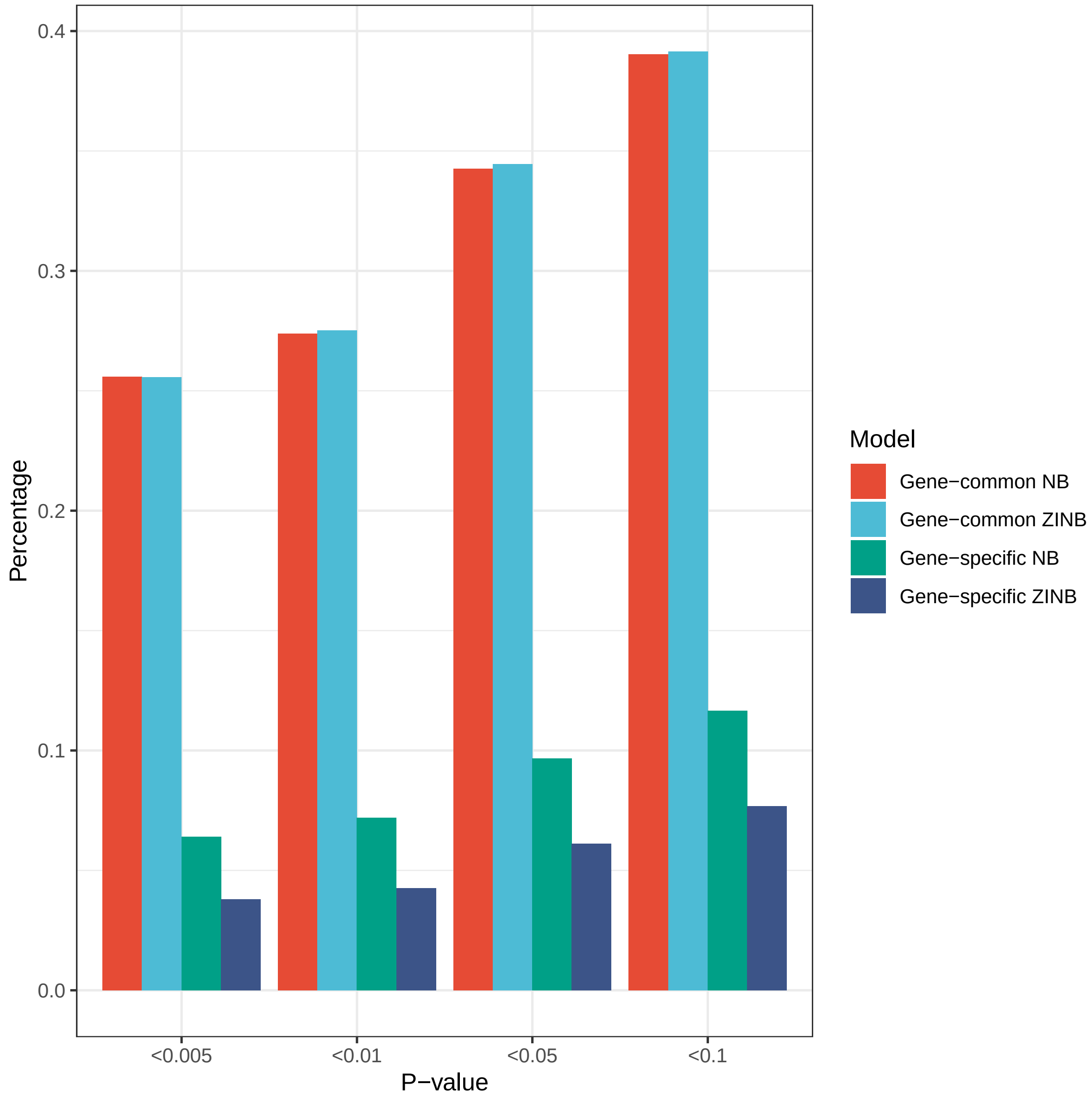


**Supplementary Figure 2: Distributions of p-values from the Pearson's $\chi^2$ goodness-of-fit test on the CT26.WT monoclonal dataset.** Tests were performed on $11,710$ genes. The x-axis is the range of p-values. The y-axis represents the proportion of genes (tests) that fall into the p-value range.

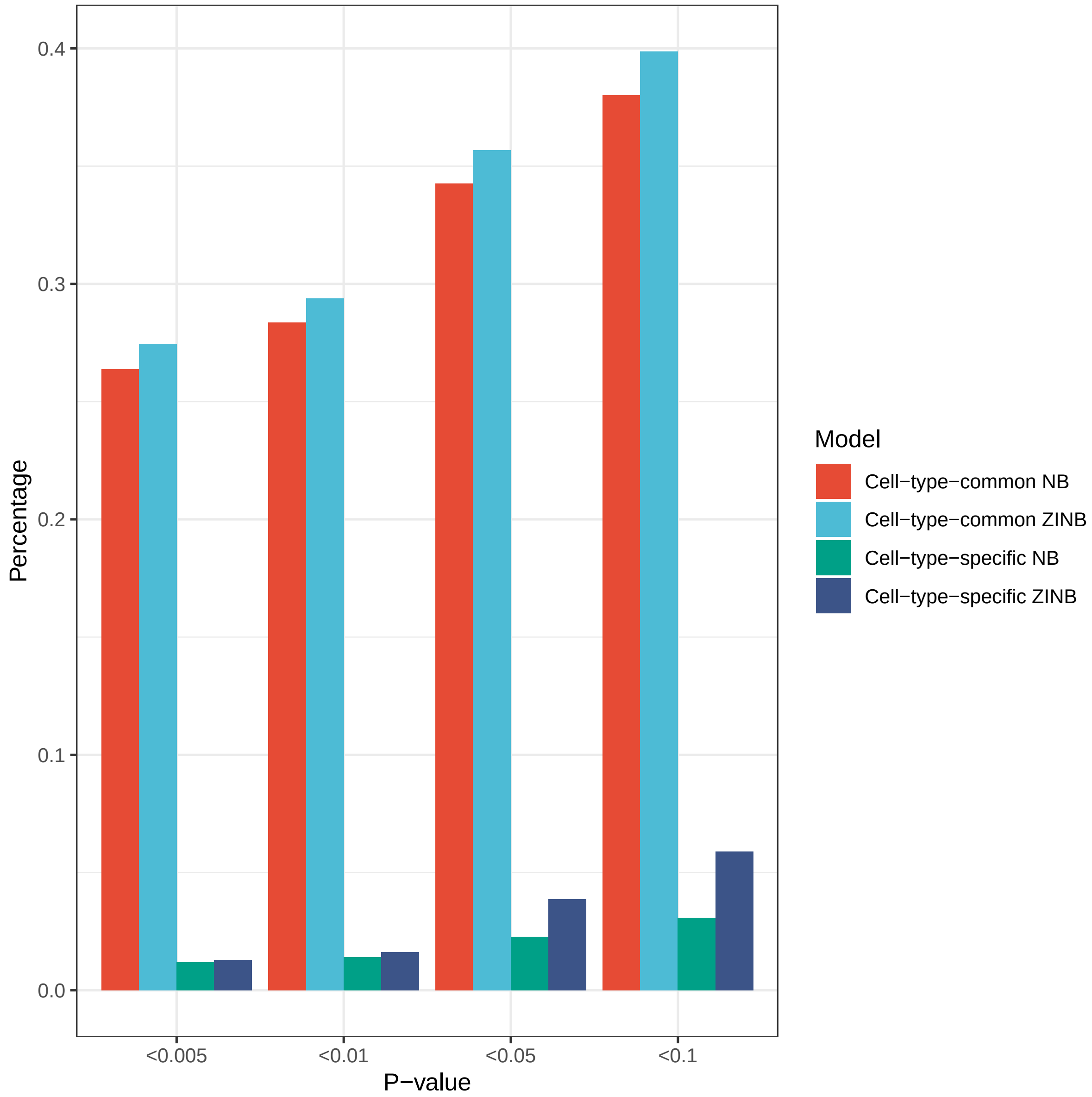


**Supplementary Figure 3: Distribution of the p-values from the cell-type-specific $\chi^2$ goodness-of-fit test on the T-CELL dataset.** Tests were performed on $7,893$ genes. The x-axis is the range of p-values. The y-axis represents the proportion of genes (tests) that fall into the p-value range.

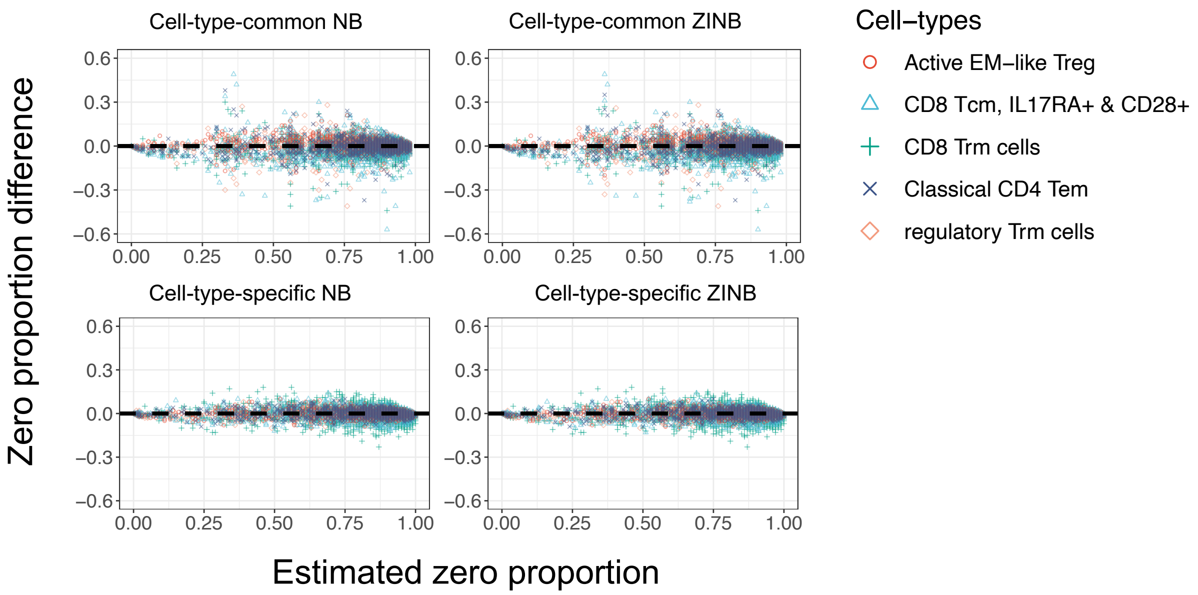


**Supplementary Figure 4: Zero UMI count proportion fitting for T-CELL.** The x-axis is the estimated zero UMI count proportion, and the y-axis is the difference between observed and the estimated zero proportions. Each point represents a gene in a specific cell-type. To reduce the size of the plot, $2,000$ genes were randomly selected for visualization.

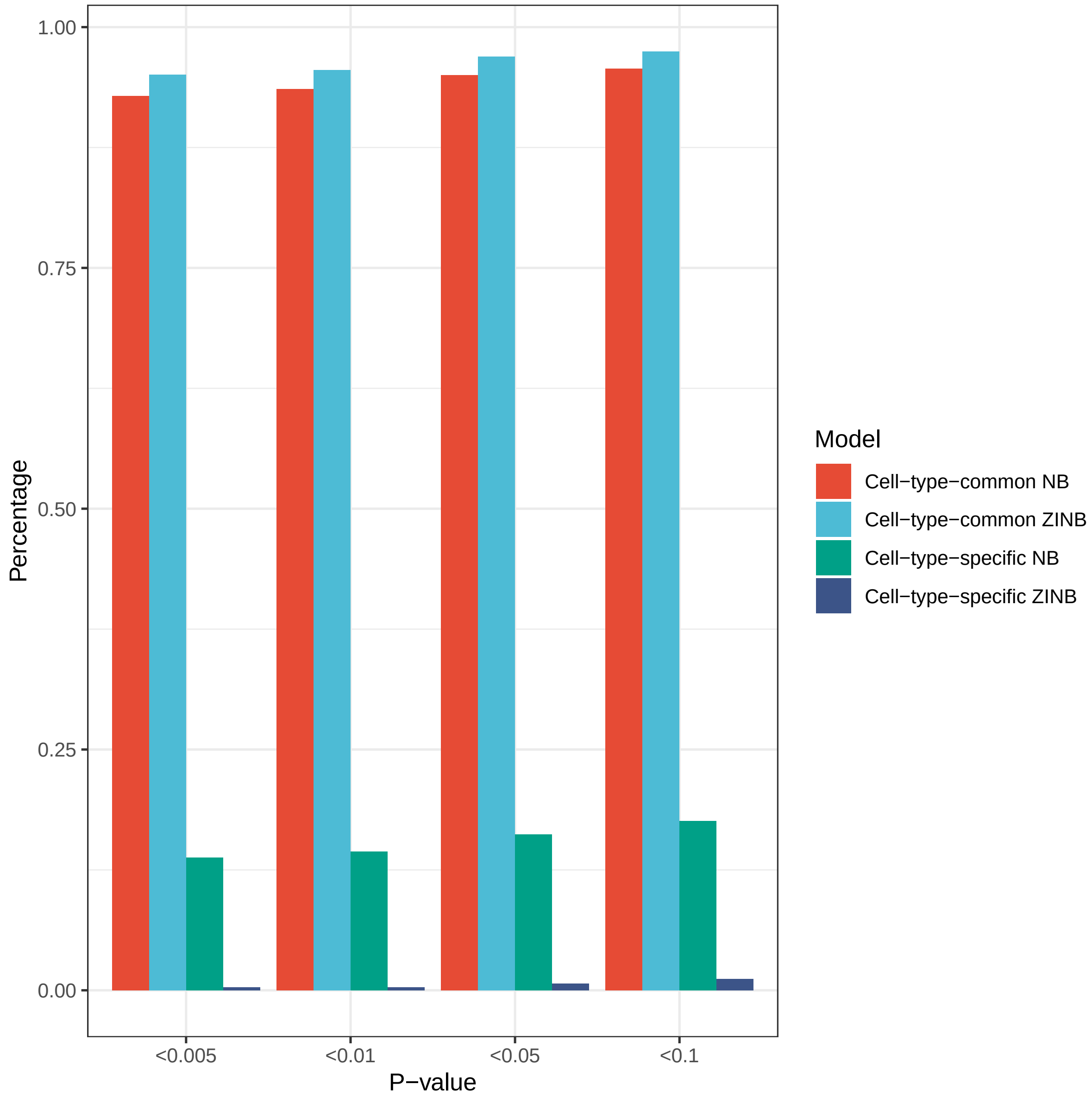


**Supplementary Figure 5: Distribution of p-values from the cell-type-specific $\chi^2$ goodness-of-fit test in CORTEX.** Tests were performed on $12,887$ genes. The x-axis is the range of p-values. The y-axis represents the proportion of genes that fall into the p-value range.

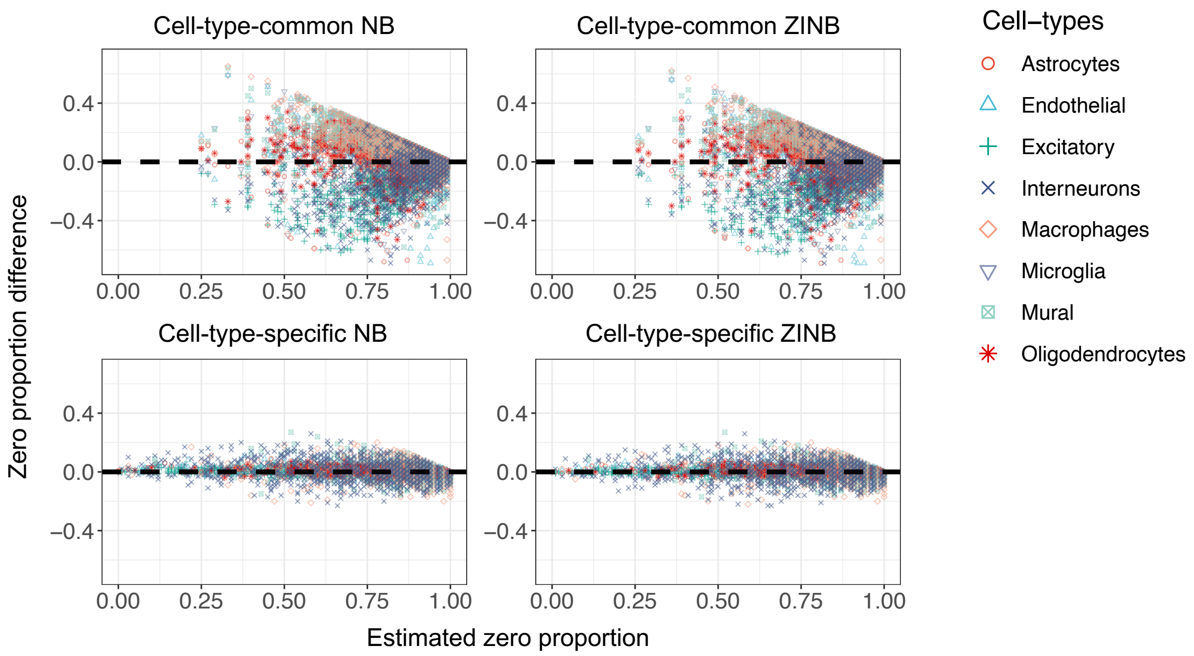


**Supplementary Figure 6: Zero UMI count proportion fitting for CORTEX.** The x-axis is the estimated zero UMI count proportion, and the y-axis is the difference between observed and the estimated proportions. Each point represents a gene in a specific cell-type. To reduce the size of the plot, 2,000 genes were randomly selected for visualization.

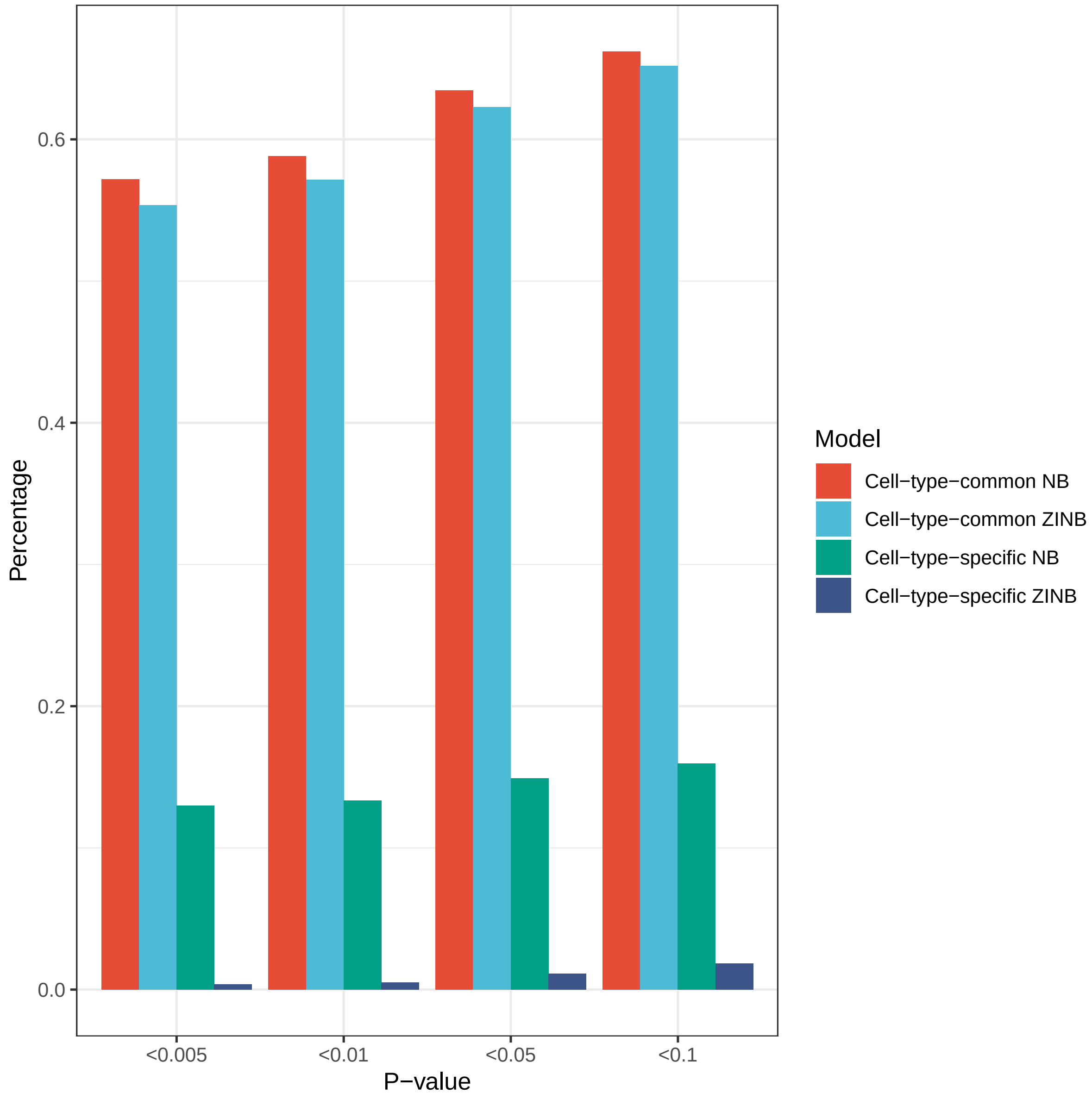


**Supplementary Figure 7: Distribution of p-values from the cell-type-specific $\chi^2$ goodness-of-fit test in RETINA Batch 1.** Tests were performed on $12,000$ genes with $13,666$ cells in Batch 1 to avoid the impact of the batch effect. The x-axis is the range of p-values. The y-axis represents the proportion of genes (tests) that fall into the p-value range. Details of test procedures can be found in Methods.

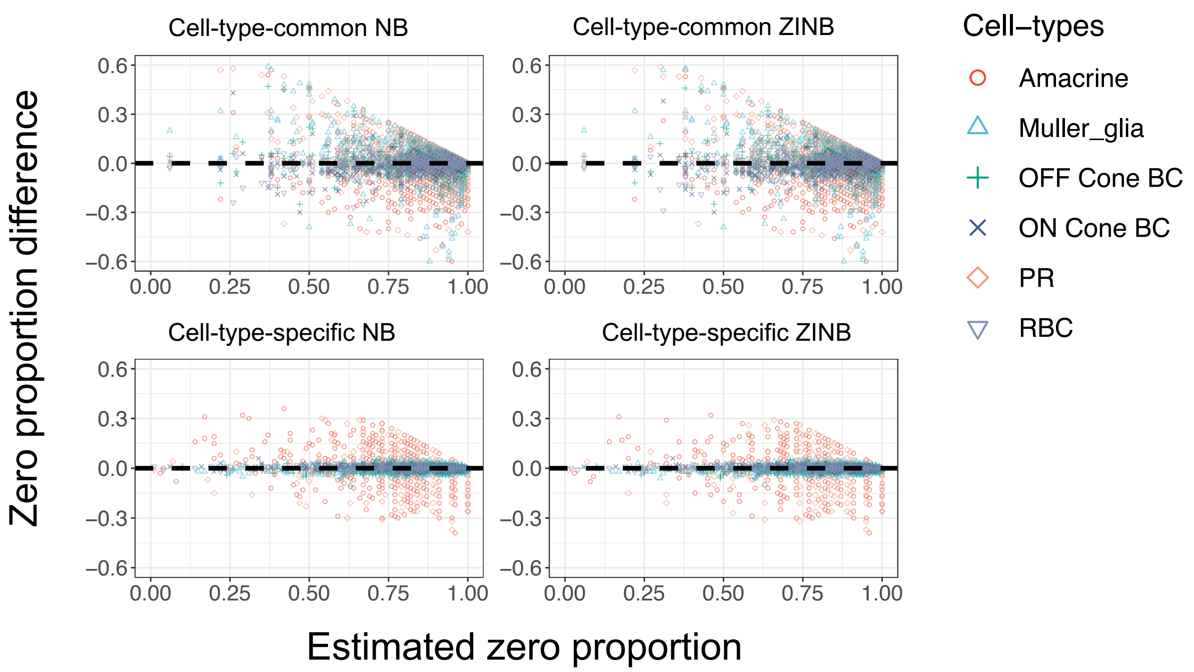


**Supplementary Figure 8: Zero UMI count proportion fitting for RETINA batch 1.** RETINA Batch 1 contains 13,666 cells, half of them are used for testing. The x-axis is the estimated zero UMI count proportion, and the y-axis is the difference between observed and the estimated proportions. Each point represents a gene in a specific cell-type. To reduce the size of the plot, 2,000 genes were randomly selected for visualization. The PR and Amacrine cell-types have relatively small number of cells (PR, 0.278%, 38 cells; Amacrine, 0.329%, 45 cells), and thus the model fittings are less accurate, especially the models are fitted using only the training samples.

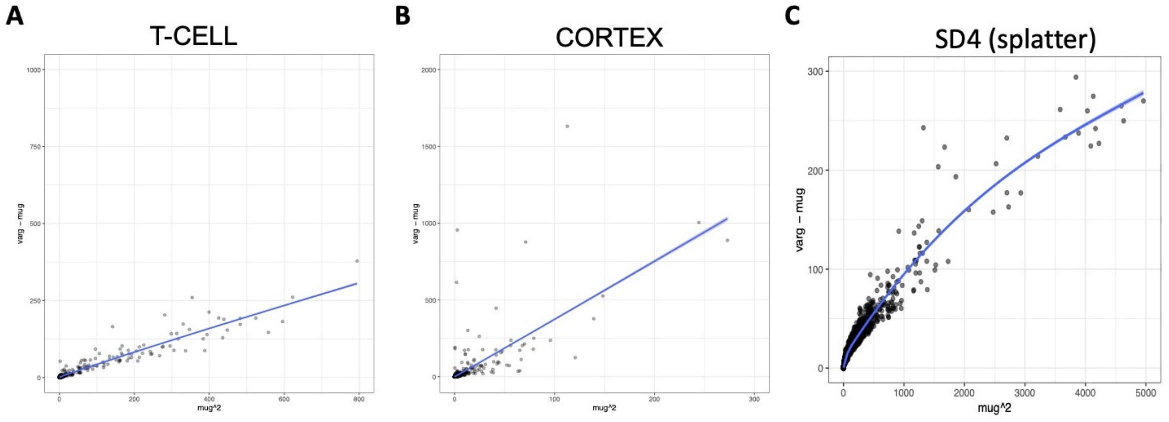


**Supplementary Figure 9: Mean-Variance trends of UMI counts in T-CELL, CORTEX, and SD4.** Each dot represents a gene. The x-axis is UMI mean square $\mu^2$; the y-axis is the UMI variance minus the UMI mean, $\sigma^2 - \mu$. The smooth curve is fitted by the generalized additive model. ($A$) T-CELL (7,893 genes). For better visualization, 6 genes with $\mu^2 \geq 800$ are not shown. ($B$)CORTEX (12,887 genes). For better visualization, 1 gene with $\mu^2 \geq 300$ is not shown. ($C$) SD4 (4,885 genes). For better visualization, 230 genes with $\mu^2 > 5000$ are not shown.

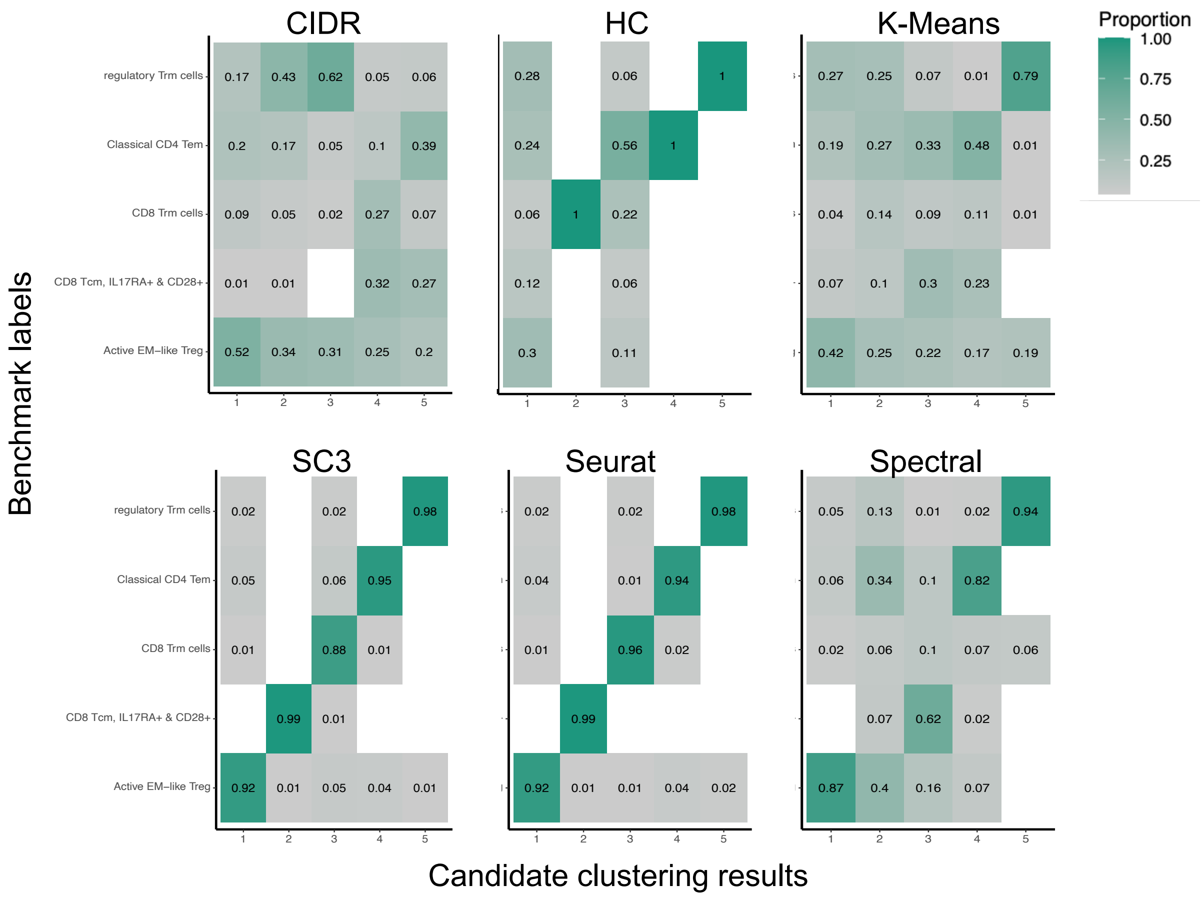


**Supplementary Figure 10: Benchmark cell proportion heatmaps.** The x-axis labels the clusters in the candidate five-cluster labels set for T-CELL, and y-axis labels the five benchmark cell types. The number and the color of each rectangle mark the benchmark cell type proportions in each candidate cluster. For example, in SC3, 92% of the cluster 1 cells are active EM-like Treg cells.

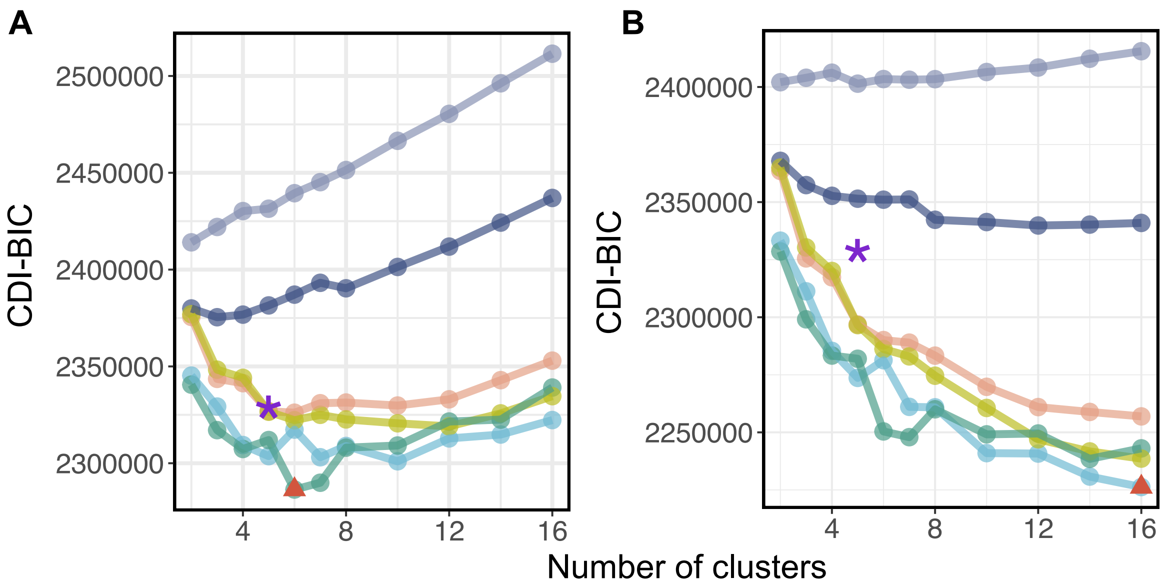


**Supplementary Figure 11: CDIs for T-CELL based on the WDS- and VST-selected feature genes.** (*A*) CDIs based on the 500 feature genes selected by WDS; (*B*) CDIs based on the 500 feature genes selected by VST. The x-axis marks the number of clusters; the y-axis marks the CDIs. Each dot represents a label set (either the benchmark or the candidate label set). The red triangle marks the CDI-selected label set. The purple star marks the benchmark label set.

**Global:**

| (GENES) | (CELLS) | [SEED] |
|---|---|---|
| 5000 | 3000 | 1 |

**Batches:**

| [BATCHES] | [BATCH CELLS] | [Location] | [Scale] |
|---|---|---|---|
| 1 | 3000 | 0.1 | 0.1 |

**Mean:**

| (Rate) | (Shape) |
|---|---|
| 0.3 | 0.6 |

**Library size:**

| (Location) | (SCALE) | (Norm) |
|---|---|---|
| 11 | 0.1 | FALSE |

**Exprs outliers:**

| (Probability) | (Location) | (Scale) |
|---|---|---|
| 0.05 | 4 | 0.5 |

**Groups:**

| [GROUPS] | [GROUP PROBS] |
|---|---|
| 5 | 0.2, 0.2, 0.2, 0.2, 0.2 |

**Diff expr:**

| [PROBABILITY] | [DOWN PROB] | [LOCATION] | [SCALE] |
|---|---|---|---|
| 0.01, 0.01, 0.01, 0.01, 0.01 | 0 | 0.4 | 0.1 |

**BCV:**

| (Common Disp) | (DoF) |
|---|---|
| 0.1 | 60 |

**Dropout:**

| [Type] | (Midpoint) | (Shape) |
|---|---|---|
| none | 0 | -1 |

**Paths:**

| [From] | [Steps] | [Skew] | [Non-linear] | [Sigma Factor] |
|---|---|---|---|---|
| 0 | 100 | 0.5 | 0.1 | 0.8 |

**Supplementary Figure 12: Parameters used in the Splatter simulator to generate SD4.**

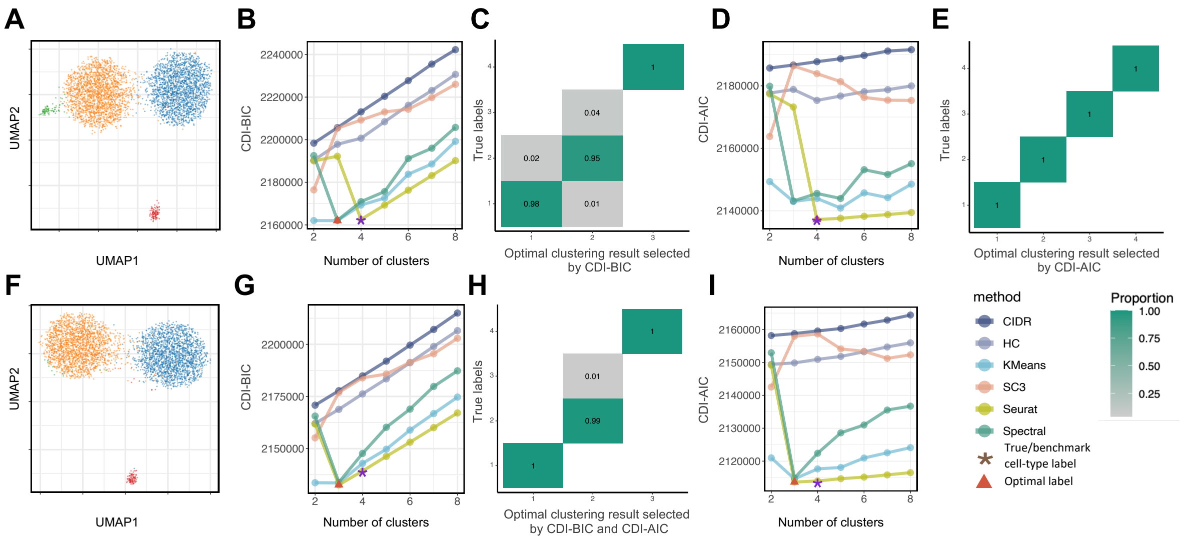


**Supplementary Figure 13: Performance of CDI on the datasets with rare cell populations with different number of cells.** $(A)-(D)$ One simulation setting with rare cell types, where each of the two normal-sized cell-types contains 2,000 cells, the rare cell type closer to the normal-sized cell types contains 85 cells, and the other rare cell type contains 100 cells; $(F)-(I)$ Another simulation setting with rare cell types, where each of the two normal-sized cell-types contains 2,000 cells, the rare cell type close to the normal-sized cell types contains 20 cells, and the other rare cell type contains 100 cells. $(A)$ and $(F)$ UMAPs based on the WDS-selected feature genes; $(B)$and $(G)$VCDI-BIC scores of the candidate and benchmark label sets; $(D)$ and $(I)$ CDI-AIC scores of the candidate and benchmark label sets. For the first simulation setting, CDI-AIC and CDI-BIC selected different optimal label sets: $(C)$ Benchmark cell type proportion heatmaps of the CDI-BIC selected label set; $(E)$ Benchmark cell type proportion heatmaps of the CDI-AIC selected label set. For the second simulation setting, CDI-AIC and CDI-BIC selected the same optimal label sets: $(H)$ Benchmark cell type proportion heatmaps of the CDI-BIC and CDI-AIC selected label set.

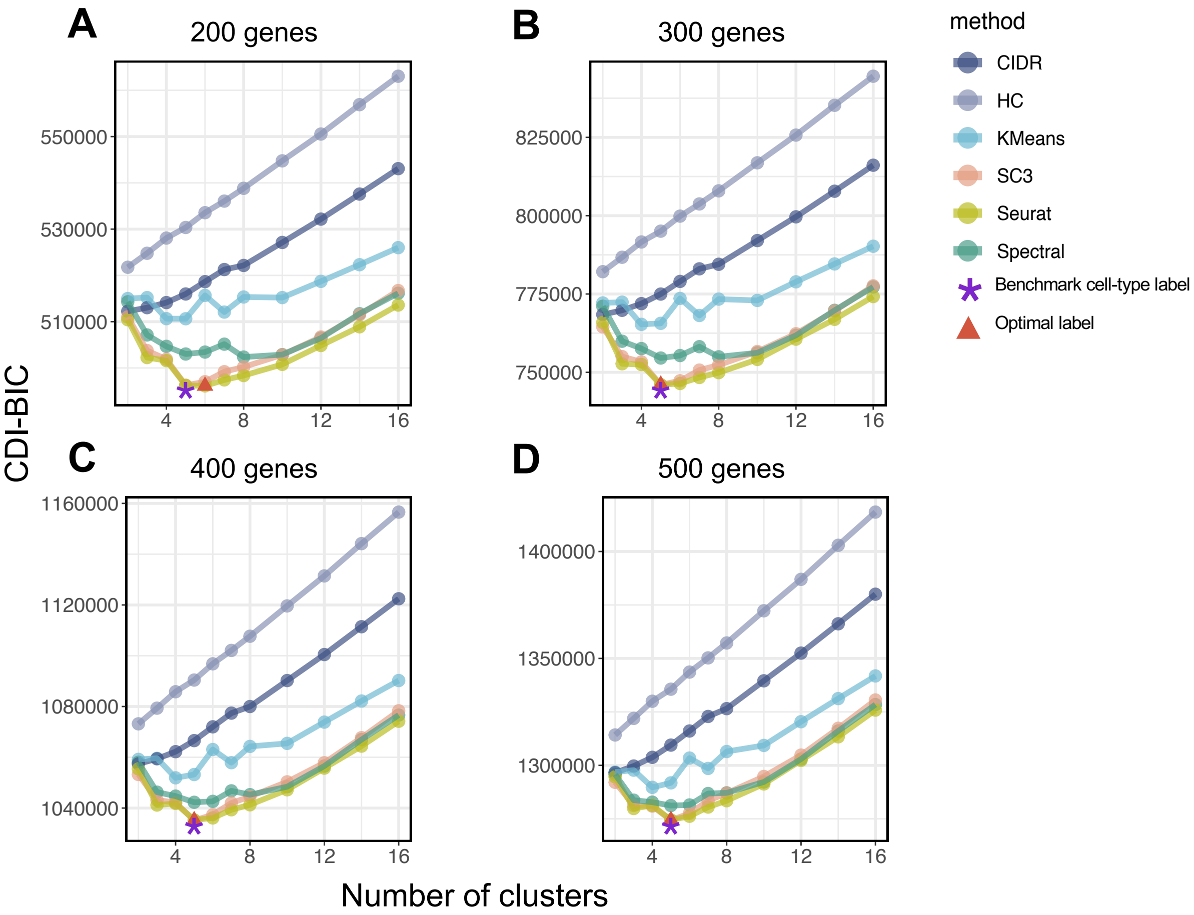


**Supplementary Figure 14: CDI performance based on different numbers of WDS-selected feature genes.** CDI-AIC was applied to the T cell dataset with different numbers of WDS-selected feature genes. (*A*) Based on the 200 WDS-selected feature genes, CDI selected the six-clsuter label set generated by Seurat. (*B*) Based on the 300 WDS-selected feature genes, CDI selected the five-cluster label set generated by Seurat. (*C*) Based on the 400 WDS-selected feature genes, CDI selected the five-cluster label set generated by SC3. (*D*) Based on the 500 WDS-selected feature genes, CDI selected the five-cluster label set generated by SC3. In all plots, the x-axis is the numbers of clusters, the y-axis is the CDI values, and the colors represent different clustering methods. The red triangle is the CDI-selected label set. The purple star denotes the benchmark label set.

### 2 Supplementary Notes

#### 2.1 Supplementary Note 1

Here we list the 8 models we considered for UMI count fitting.

##### Notations

- $c$: index of cell ($c = 1, \ldots, N$)
- $g$: index of gene ($g = 1, \ldots, G$)
- $k$: index of cell-type ($k = 1, \ldots, K$)
- $X_{gc}$: raw UMI count for gene $g$ in cell $c$
- $s_c$: size factor for cell $c$
- $\phi$: dispersion parameter shared by all genes and all cells in one dataset
- $\delta$: zero inflation factor shared by all genes and all cells in one dataset
- $\mu_g$: mean parameter for gene $g$ across all cells
- $\phi_g$: dispersion parameter for gene $g$ across all cells
- $\delta_g$: zero-inflation factor for gene g across all cells
- $\mu_{g,k}$: mean parameter for gene g in cell-type $k$
- $\phi_{g,k}$: dispersion parameter for gene $g$ in cell-type $k$
- $\delta_{g,k}$: zero-inflation factor for gene $g$ in cell-type $k$

##### Model distributions

In general, if $Y \sim \text{NB}(\mu, \phi)$, then the probability mass function of $Y$ is

$$P(Y = y) = F(y \mid \mu, \phi) = \frac{\Gamma(y + 1/\phi)}{\Gamma(y + 1)\Gamma(1/\phi)} \left(\frac{1/\phi}{\mu + 1/\phi}\right)^{1/\phi} \left(\frac{\mu}{\mu + 1/\phi}\right)^{y}, \forall y \in \mathbb{N} \cup \{0\}. \tag{1}$$

In the manuscript, the models we considered are listed in Supplementary Table **??**.

| Model name | Distribution of $X_{gc}$ |
|---|---|
| Gene-common NB | $\mathrm{NB}(s_c\mu_g, \phi)$ |
| Gene-common ZINB | $\mathrm{ZINB}(s_c\mu_g, \phi, \delta)$ |
| Gene-specific NB | $\mathrm{NB}(s_c\mu_g, \phi_g)$ |
| Gene-specific ZINB | $\mathrm{ZINB}(s_c\mu_g, \phi_g, \delta_g)$ |
| Cell-type-common NB | $\mathrm{NB}(s_c\mu_g, \phi_g)$ |
| Cell-type-common ZINB | $\mathrm{ZINB}(s_c\mu_g, \phi_g, \delta_g)$ |
| Cell-type-specific NB | $\mathrm{NB}(s_c\mu_{g,k}, \phi_{g,k})$ for cell $c$ in cell-type $k$ |
| Cell-type-specific ZINB | $\mathrm{ZINB}(s_c\mu_{g,k}, \phi_{g,k}, \delta_{g,k})$ for cell $c$ in cell-type $k$ |

**Supplementary Table 1: Model names and the corresponding distributions of raw UMI count.**

##### Size factor estimation

The size factor estimation follows the DESeq2 median-of-ratios procedure. The size factor of cell $c$ is estimated with

$$\hat{s}_c = \mathrm{median}_g \frac{\max\{X_{gc}, 0.5\}}{\left(\prod_{c=1}^{N} \max\{X_{gc}, 0.5\}\right)^{1/N}}.$$

We add 0.5 to zero count to avoid the zero denominators.

#### 2.2 Supplementary Note 2

For simplicity we omit the batch index $b$ in the superscript (if any) and size factor $s_c$. Suppose

$$L_{0,c} \sim \mathrm{Multinomial}(K_0; \pi_1, \dots, \pi_{K_0}).$$

Combining with the NB model

$$X_{gc} \mid (L_{0,c} = k) \sim \mathrm{NB}(\mu_{gk}, \phi_{gk}),$$

we have $X_{gc} \sim \sum_{k=1}^{K_0} \pi_k \mathrm{NB}(\mu_{gk}, \phi_{gk})$ with

$$\xi_{1,g} = \mu_g = \mathtt{E}(X_{gc}) = \sum_{k=1}^{K_0} \pi_k \mu_{gk}, \quad \xi_{2,g} = \mu_g^2 + \sigma_g^2 = \mathtt{E}(X_{gc}^2) = \sum_{k=1}^{K_0} \pi_k \{\mu_{gk} + (1+\phi_{gk})\mu_{gk}^2\}.$$

Therefore, $\xi_{2,g} - \xi_{1,g} = \sum_{k=1}^{K_0} \pi_k (1+\phi_{gk})\mu_{gk}^2$. Let $\xi_{3,g} = \sum_{k=1}^{K_0} \pi_k (1+\phi_{gk})^{1/2}\mu_{gk}$. Then

$$\eta_g = \frac{\xi_{2,g} - \xi_{1,g}}{\xi_{3,g}^2} = \sum_{k=1}^{K_0} \pi_k \left\{ \frac{(1+\phi_{gk})^{1/2}\mu_{gk}}{\xi_{3,g}} \right\}^2.$$

Let

$$a_k = \pi_k^{1/2} \frac{(1+\phi_{gk})^{1/2}\mu_{gk}}{\xi_{3,g}}, \quad b_k = \pi_k^{1/2}.$$

By $a_k^2 \geq 2a_k b_k - b_k^2$, we have

$$\eta_g = \sum_{k=1}^{K_0} a_k^2 \geq 2 \sum_{k=1}^{K_0} a_k b_k - \sum_{k=1}^{K_0} b_k^2 = 2 \sum_{k=1}^{K_0} \frac{\pi_k (1+\phi_{gk})^{1/2} \mu_{gk}}{\xi_{3,g}} - \sum_{k=1}^{K_0} \pi_k = 1.$$

The minimum is obtained when $a_k = b_k$ for all $k \in \{1, \ldots, K_0\}$, *i.e.*,

$$(1+\phi_{gk})\mu_{gk}^2 = C_g, \tag{2}$$

not depending on the cell type $k$.

If gene $g$ is not a feature gene, then $\mu_{gk} = \mu_g$ and $\phi_{gk} = \phi_g$, thus (**??**) holds, and $\eta_g$ will reach the minimum 1.

If gene $g$ is a feature gene, then as long as $(1+\phi_{gk})\mu_{gk}^2$ are not all equal across cell type $k$, $\eta_g > 1$. Thus for feature genes, $\eta_g$ tends to be large.

Unfortunately, $\xi_{3,g}$ is hard to be estimated if the true cell labels are unknown. Thus, $\eta_g$ is hard to be estimated directly. We need to find a surrogate $\xi_g$ to approximate $\eta_g$.

It is easy to see that the WDS for gene $g$ is

$$\xi_g = \frac{\sigma_g^2 - \mu_g}{\mu_g^2} = \frac{\xi_{2,g} - \xi_{1,g}^2 - \xi_{1,g}}{\xi_{1,g}^2} = \frac{\xi_{2,g} - \xi_{1,g}}{\xi_{1,g}^2} - 1.$$

When $\phi_{gk} \approx \phi$ for all $g$ and $k$, $\xi_{3,g} \approx (1+\phi)^{1/2}\xi_{1,g}$. Then, $\xi_g \approx \eta_g(1+\phi) - 1$.

In general, let $\phi_{\max} = \max_{k,g} \phi_{g,k}$. Then $\eta_g$ can be bounded on both sides:

$$\frac{1}{1+\phi_{\max}}(\xi_g + 1) \leq \eta_g \leq \xi_g + 1.$$

We assume that $\phi_{\max}$ are not too large. Then $\xi_g$ can be used as a surrogate of $\eta_g$. We estimate $\xi_g$ by

$$\hat{\xi}_g = \frac{\hat{\sigma}_g^2 - \hat{\mu}_g}{\hat{\mu}_g^2}.$$
